# Disentangling the Effects of Intercropping on Vector-Borne Plant Virus Dynamics

**DOI:** 10.1101/2025.07.17.664952

**Authors:** Blake Corrigan, Jon Yearsley

## Abstract

Emerging plant diseases threaten crop yields, food security and the health of ecosystems. In fields with multiple plant species, disease management can become challenging, as different host plants can either reduce or increase virus transmission. Motivated by empirical evidence that intercropping tends to limit virus spread, we develop a mathematical model that integrates the epidemiological dynamics of multiple plants and a vector population. We derive the basic reproduction number and quantify how relative differences in intercrop characteristics determines outbreak persistence. We find conditions when intercropping reduces the average number of secondary infections from a plant compared to a monoculture. We also demonstrate cases where intercropping can increase the risk of a disease outbreaks. Crucially, even when disease persistence is unlikely, transient outbreaks may still occur. We investigate such short-term dynamics by deriving a threshold index that predicts when transient outbreaks are possible. We give conditions that ensure there are plant mixture compositions where persistent and transient outbreaks are unlikely in a monoculture and/or inter-cropped system. We discuss the practical implications of all our results, which provide a theoretical foundation for sustainable plant disease control.

## 1 Introduction

Vector-borne plant viruses are a major concern in agronomy, as they often lead to substantial yield losses and economic damage (Ristaino et al., 2021). Controlling diseases caused by such viruses remains challenging due to the complex interactions among viruses, plant hosts, and insect vectors (Hosack et al., 2008). While insecticides are often used to reduce vector population sizes, their effectiveness can be limited by rapid reinvasion, vector resistance, and negative environmental impacts (Gong et al., 2023). Intercropping is the simultaneous cultivation of multiple crop species within the same field. This agricultural practice has emerged as a sustainable strategy that can not only produce higher yields than monocultures, but mitigate virus spread, strengthen agroecosystem resilience and reduce susceptibility to pest invasions (Grauby et al., 2022; Hooks & Fereres, 2006; O’Malley et al., 2025; Roudine et al., 2025; Tous-Fandos et al., 2025).

Building on such empirical findings, mathematical modelling offers a complementary quantitative framework that can deepen our understanding of how intercrop traits and management practices influence disease dynamics. Recent theoretical models of plant virus dynamics have incorporated spatial structure to assess how crop arrangements influence disease spread (Allen-Perkins & Estrada, 2019; Rosales Herrera et al., 2021; Rother et al., 2025); have focused on specific crop–vector–virus systems to evaluate species-targeted interventions (Cunniffe et al., 2021; Falla & Cunniffe, 2024); or have examined how resistant varieties affect both disease dynamics and crop yield (Vyska et al., 2016). Although these models provide valuable insights into various aspects of vector-mediated plant-virus dynamics, they often consider only a single crop species or do not derive general analytical expressions to guide disease management. We present a theoretical framework that tracks virus transmission between multiple host crops and an insect vector, enabling us to capture essential transmission dynamics and derive explicit expressions for key indicators related to outbreak risks.

Our intercropping model builds upon the eco-epidemic frameworks of Chamchod and Britton (2011), Cunniffe et al. (2021), Falla and Cunniffe (2024), Hamelin et al. (2023), Madden et al. (2000), and Roosien et al. (2013), which have highlighted key mechanisms that drive vector-borne disease spread within single-host systems. It is worth noting that in our multi-host framework, instead of each crop being a different species, each could be a variety of the same crop species, such as resistant or non-resistant types (Murray-Watson & Cunniffe, 2022). In intercropping systems, the intercrop could also be a so-called sentinel plant, which is a highly susceptible host species that displays visible infection symptoms quicker than the main crop (Lovell-Read et al., 2023). In this scenario, for example, the virus may be diluted within the host community, but the infected sentinel plants may also be able to be removed rapidly enough, with both of these limiting of disease spread. Using a multi-host modelling framework can be useful not only to capture the dynamics of multiple cultivated species, but also to model systems where viral hosts include weeds or plants within field margins (Roudine et al., 2025; Weber et al., 2026). These are important examples to consider, as unmanaged hosts such as weeds could sustain vectors outside of growing seasons, acting as entry points for vectors once the growing season begins (Yazdkhasti et al., 2021).

Examples of virus-host-vector systems that our model could capture include yellow dwarf viruses, persistent viruses transmitted by several aphid species that infects multiple cereal crops, such as rice, barley, oat and maize (McElhany et al., 1995); cassava mosaic geminiviruses, a group of related virus species that infect cassava, castor bean and cotton crops, and are vectored by whiteflies (Legg et al., 2014); tomato yellow leaf curl virus, a persistent begomovirus which infects tomato, pepper and bean crops worldwide and is vectored by whiteflies (Riley & Srinivasan, 2019); cucumber mosaic virus, a non-persistent virus vectored by many aphid species, with a broad host range which includes cucumber, squash and pepper crops (Gallitelli, 2000); and potato virus Y, a potyvirus transmitted by aphids, that infects multiple hosts, including potato, tobacco and pepper crops (Kaliciak & Syller, 2009).

Although our model is framed in terms of plant viruses, the general structure could also be applied to bacterial or fungal pathogens that are vectored between plants. These could include *Xylella fastidiosa*, a bacterial pathogen spread by xylem sap-feeding insects, such as spittlebugs, causing severe damage to a wide range of plant species, such as lavender, rosemary, olive and coffee crops (Cornara et al., 2019); and some *Ophiostomatales* fungal pathogens vectored by bark beetles, such as *Ips typographus*, that infect several conifer species (Kandasamy et al., 2023). The pathogens in all of these example are mainly horizontally transmitted between vectors and plants.

The number of host plants in a field plays a critical role in reducing or amplifying the severity of disease outbreaks. The dilution effect hypothesis proposes that increasing host diversity often reduces disease risk by interrupting transmission pathways (Keesing & Ostfeld, 2021). In contrast, the amplification effect occurs when greater host diversity leads to increased pathogen spread, by raising overall host densities for example. As described by McCann and Gellner (2020), whether dilution or amplification dominates within a system depends on factors such as host competence, community composition, and the mode of transmission.

This complexity highlights why it is important to study the effects of intercropping when predicting how biodiversity influences outbreak risk within heterogeneous environments.

We present our theoretical framework by using it to answer three applied questions:

Q1 Which intercropping strategies most effectively limit the risk of a persistent plant-virus outbreak?

Q2 In what ways does intercropping alter persistent outbreak risk comapared to monocultures?

Q3 When persistent outbreaks are unlikely, how does intercropping shape transient outbreak potential?

To answer Q1 we first derive the basic reproduction number (a persistent outbreak indicator) and then explore how this quantity behaves under intercrop parameter variation. We answer Q2 by comparing how the basic reproduction number differs between monoculture vs intercropping systems. Lastly, to answer Q3 we derive the threshold index of epidemicity. After we answer each of these questions, we discuss the ecological consequences of the results and how these may correspond to practical management strategies in each case.

## 2 The Multi-Host Model

Our model extends the base models of Chamchod and Britton (2011), Falla and Cunniffe (2024), Hamelin et al. (2023), and Roosien et al. (2013). In each of these papers the authors investigate a particular single-host model of vector-borne pathogen transmission. We use a similar Susceptible-Infected approach to model pathogen transmission dynamics in a single-vector and multi-crop community. The field system comprises *n* ≥ 2 distinct crop species. An individual crop can be either infected or susceptible (i.e. not infected), where *P*_*k,I*_ and *P*_*k,S*_ are respectively the number of infected and susceptible individuals for crop *k*. The vector population is likewise divided into the number of susceptible (*V*_*S*_) and the number of infected (*V*_*I*_) individuals.

To make interpretation of our model more direct, we designate species *k* = 1 as a primary crop species of interest, typically a high-value agricultural plant. The remaining species, indexed by *k* ∈ {2, …, *n*}, represent various intercrops or surrounding non-crop vegetation. These intercrops may differ in their epidemiological roles, acting as reservoirs or barriers to transmission depending on their infectivity and attractiveness properties.

The dynamics of this system is governed by

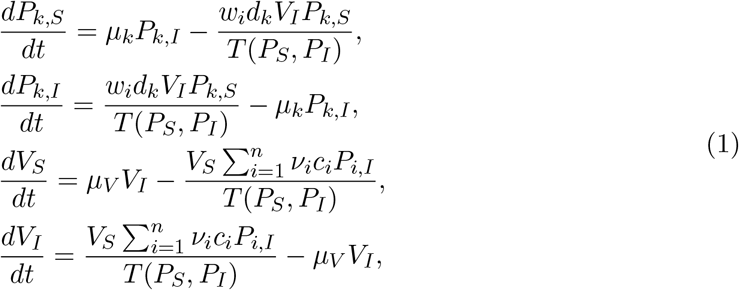

for *k* ∈ {1, …, *n*}. The interpretations of parameters in (1) are given in Table 1. Within our model we are ignoring spatial structure and so assume that the arrangement of crop species does not alter the dynamics between susceptible and infected compartments. From the form of 1 we can also see that each crop and vector population has constant total population sizes, i.e. *P*_*k,S*_(*t*) + *P*_*k,I*_(*t*) = *P*_*k*_ and *V*_*S*_(*t*) + *V*_*I*_(*t*) = *V*_*T*_ for all *t* ≥ 0, for constants *P*_*k*_, *V*_*T*_ ≥ 0.

**Table 1:**
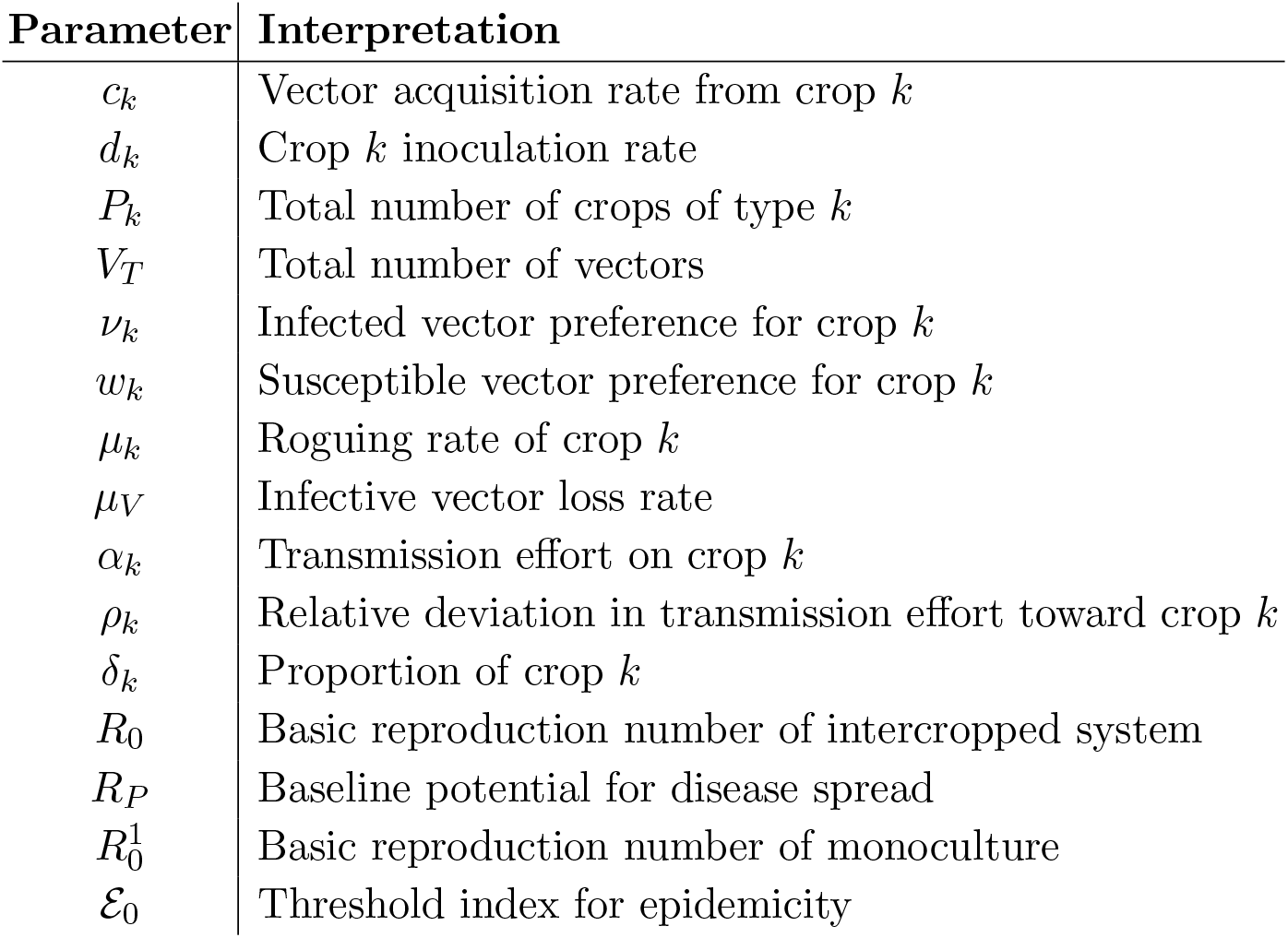
Glossary of parameters within (or derived from) the model and their interpretation.

Virus vectors often respond to specific cues, which can be visual, olfactory, or otherwise, and this can vary with plant infection status (Chamchod & Britton, 2011; Falla & Cunniffe, 2024; Hamelin et al., 2023). With this in mind, we assume that vectors can be biased either in favour of (*ν*_*k*_ *>* 1), or against (0 ≤ *ν*_*k*_ *<* 1), landing on an infected crop of type *k*. We also assume that vectors can be biased either in favour of (*w*_*k*_ *>* 1), or against (0 ≤ *w*_*k*_ *<* 1), landing on a susceptible crop of type *k*. Note that *w*_*k*_ = 1 (*ν*_*k*_ = 1) corresponds to the vector having no preference for the *k*th susceptible (infected) crop. Let *P*_*S*_ := (*P*_1,*S*_, …, *P*_*n,S*_)^*T*^, *P*_*I*_ := (*P*_1,*I*_, …, *P*_*n,I*_)^*T*^. Given that we have *n* crops, we assume that the probability of vectors landing on the *kth* susceptible crop is

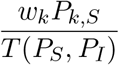

and the probability of vectors landing on the *kth* infected crop is

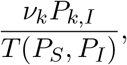

where

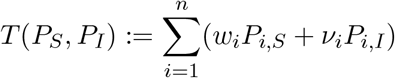

controls how vectors allocate their probing or feeding effort across the host community. One can verify that these probabilities sum to unity over all *n* plants, and so are each well-defined.

The rate at which susceptible vectors acquire the virus from infected crop *k* is controlled by the parameter *c*_*k*_ ≥ 0. The rate at which a susceptible crop *k* acquires the virus from an infected vector is controlled by the parameter *d*_*k*_ ≥ 0. We interpret the parameter *µ*_*k*_ ≥ 0 as the rate at which infected crops are removed from the system. As total population levels are kept constant, *µ*_*k*_ is also the rate at which infected crops are replenished once removed, which then enter the susceptible compartment. This removal and replenishment of infected crops is known as roguing (Falla & Cunniffe, 2024).

Cunniffe et al. (2021) and Falla and Cunniffe (2024) interpret *µ*_*V*_ ≥ 0 as the rate at which a virus vector loses infectivity. This is particularly relevant for so-called non-persistent viruses. Another interpretation, and one that is relevant for persistent viruses, is that this may also be interpreted as the mortality rate of infected vectors, which are in turn replenished as susceptibles. This argument is analogous to the interpretation of *µ*_*k*_ and is justified based on the fact that the total population is constant and our assumption that there is no vertical transmission of the virus. Within our framework we assume that there is no vertical transmission of the virus from parent to offspring.

## 3 Threshold Conditions for Persistent Plant Virus Outbreaks

A central focus in epidemiological modelling is determining when a disease will successfully invade a population. This can be measured mathematically using the basic reproduction number, denoted by *R*_0_, which quantifies the average number of secondary infections caused by a single infected individual in an otherwise susceptible population (Van den Driessche, 2017).

It is important to show that biological models have positive solutions. In the Supplementary Material we show that this is the case for our model. Since the total populations are constant, we then have that

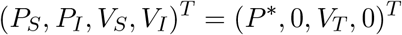

is the so-called disease-free equilibrium, the steady state of (1) where there are no infected compartments present, where *P*\* = (*P*_1_, …, *P*_*n*_)^*T*^.

Given that each total plant population remains constant over time, it suffices to focus on the infected compartments when analysing our model, since we can write *P*_*k,S*_ = *P*_*k*_ − *P*_*k,I*_ and *V*_*S*_ = *V*_*T*_ − *V*_*I*_. Thus, we can reduce our system (1) to

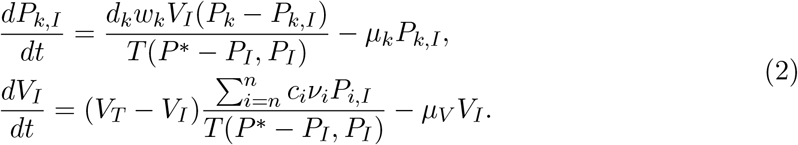

The dynamics of (2) must stay within realistic bounds. In the Supplementary Material we show that compartment trajectories do not exceed their respective total population sizes.

In order to characterise local stability of the disease-free equilibrium of (2), we computed the basic reproduction number using the next-generation matrix approach (Van den Driessche & Watmough, 2002) as

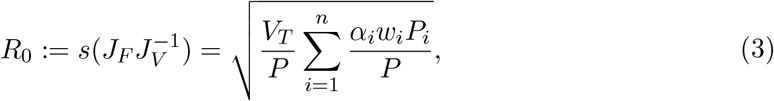

where *s*(*A*) is the spectral radius of a matrix *A* (maximum modulus over all eigenvalues). The next-generation matrix, 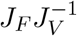, and derivation of *R*_0_ is given in the Supplementary Material. The parameter

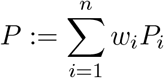

is the total weighted plant density/abundance across all hosts. The parameters *w*_*i*_ and *P*_*i*_ always appear as the product, *w*_*i*_*P*_*i*_, within our main results. If *w*_*i*_*P*_*i*_ is large, then this implies that crop type *i* is abundant and/or the susceptible sub-population of this crop type is highly attractive, and so vectors are more likely to visit, feed and infect such susceptible plants. The parameter

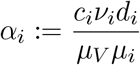

represents the virus transmission effort for plant *i*, assuming that *µ*_*V*_, *µ*_*i*_ ≠ 0. Specifically, *α*_*i*_ is the product of the rate of vector infection relative to the infected vector loss and the rate of plant infection relative to infected plant loss.

It is well known (see Supplementary Material) that if *R*_0_ *<* 1, then the disease-free equilibrium is locally asymptotically stable, meaning all sufficiently small initial infected densities of (1) result in infected trajectories decaying to 0 over time. Conversely, if *R*_0_ *>* 1, the disease-free equilibrium is unstable, and infected trajectories of (1) diverge from 0 in the long run. Throughout this paper, unless stated otherwise, the term “*persistent outbreak*” refers to *R*_0_ *>* 1, whereas “*transient outbreak*” refers to the phenomenon when the number of infected plants have the potential to initially increase even though *R*_0_ *<* 1 (details of which are given in a later section). We conducted several numerical simulations and found that in all observed scenarios, *R*_0_ *<* 1 resulted in disease extinction for all chosen positive initial conditions, i.e. this may in fact be sufficient for global asymptotic stability of the disease-free equilibrium (see the Supplementary Material for more on this).

Within a monoculture, for vector-borne diseases, the basic reproduction number, 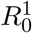, is inversely proportional to host density, since vectors are assumed to make a constant number of host visits per unit time, independent of host abundance (Falla & Cunniffe, 2024). This relationship arises from (3) as

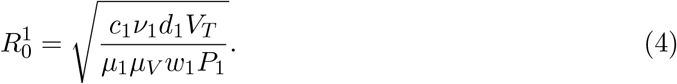

This formula clearly shows that decreasing (increasing) the density of a single host has the potential to concentrate (dilute) the virus within a population (Wonham et al., 2006). Our approach furthers this, as seen in (3), by allowing dilution/concentration effects to take place due to the presence of additional hosts, the consequences of which are quite complex.

## 4 Intercrop Characteristics and Persistent Outbreak Potential

In this section, we provide some insight into answering Q1 and explore how different parameters influence persistent disease outbreaks via *R*_0_. We can observe from (3) that in order to ensure a persistent outbreak does not occur for sufficiently small initial infected densities, i.e. *R*_0_ *<* 1, it is necessary that

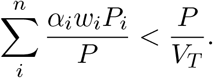

To avoid a persistent outbreak, the average infection gain rate must not exceed the critical threshold given by the weighted plant-to-vector ratio. In a monoculture system, controlling the epidemic requires *α*_1_ *< w*_1_*P*_1_*/V*_*T*_. With intercropping, however, the overall persistent outbreak risk depends on the weighted average infectivity, which is determined by both the total crop population sizes and the viral transmission efforts on each crop. This means that intercropping, with a less susceptible crop can, in general, effectively reduce *R*_0_ below the epidemic threshold even if in a monoculture there is a persistent outbreak. The parameters *α*_1_, …, *α*_*n*_ represent the virus’s transmission effort among crops and are key in determining whether an epidemic can be sustained. For the *n* = 2 case (Fig. 1), the threshold curve *R*_0_ = 1 in the (*α*_1_, *α*_2_)-plane separates regions where *R*_0_ *<* 1 and *R*_0_ *>* 1. We can see from Fig. 1 that higher transmission effort on one plant must be balanced by lower effort on the other to remain below *R*_0_ = 1.

**Fig. 1:**
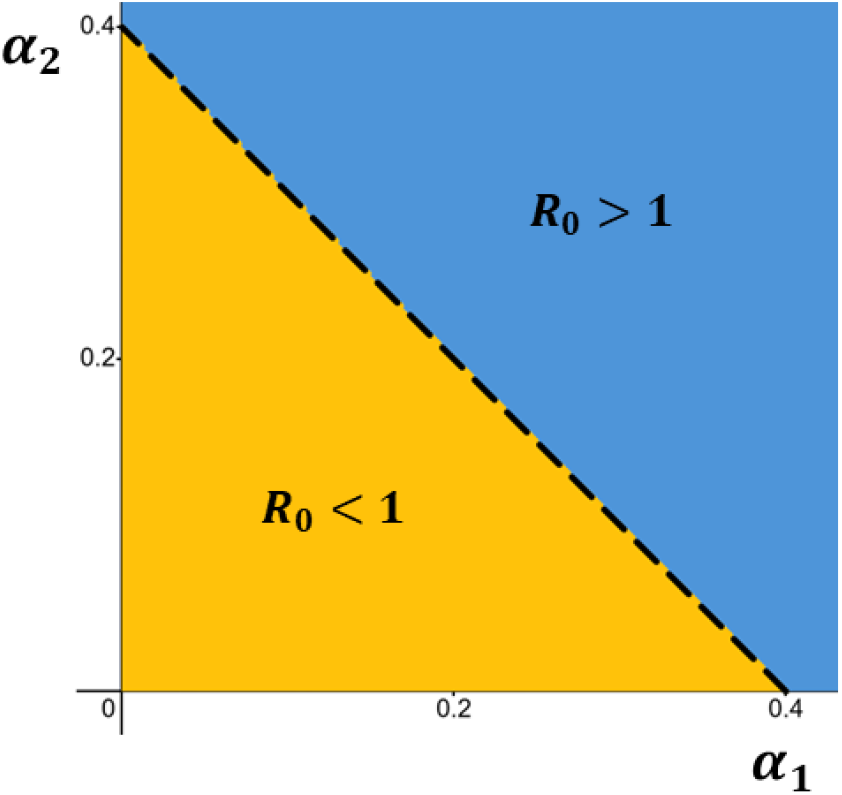
The (*α*_1_, *α*_2_)-plane showing regions where persistent outbreaks (blue) and a stable disease-free equilibrium (yellow) occur. The dashed black line is *R*_0_ = 1. Parameters: *P*_1_ = 50, *P*_2_ = 200, *V*_*T*_ = 1000, *w*_1_ = 2, *w*_2_ = 0.5.

### 4.1 Relative Deviations in Viral Transmission Effort

To better understand the behaviour of *R*_0_ as we vary crop and intercrop parameters, let

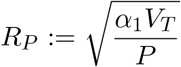

be baseline for disease spread when *P* plants are present. Define *δ*_*k*_ = *w*_*k*_*P*_*k*_*/P* as the weighted proportion of plant type *k* in the system, and

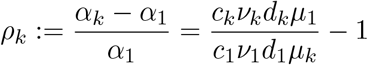

as the relative deviation in viral transmission effort for crop *k*, adjusted for loss, compared to the crop plant, respectively (see Table 1). Since *ρ*_1_ = 0, the basic reproduction number can be written compactly as

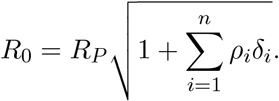

Note that when all plant types have the same infectivity and roguing parameters as the crop (*ρ*_*i*_ = 0 for all *i* ≥ 2), we recover *R*_0_ = *R*_*P*_. The baseline *R*_*P*_ provides a useful benchmark. If it is large, the system may already be conducive to epidemic spread due to factors like high vector density, high transmission efficiency from the crop, or low crop removal rate. Conversely, a small *R*_*P*_ indicates low transmission potential from the crop alone. We can easily observe that

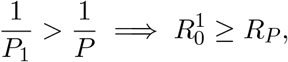

with equality only when *P* = *w*_1_*P*_1_. This highlights that introducing additional plant types may dilute the crop’s contribution to disease spread.

To better understand and visualise how *ρ*_*i*_ and *δ*_*i*_ alter *R*_0_, we will focus on the *n* = 2 case. Throughout the rest of this paper, when *n* = 2, we will write *ρ* := *ρ*_2_ and *δ* := *δ*_2_, for ease of exposition. The influence of *R*_*P*_ on *R*_0_ is illustrated in Fig. 2, showing how differences in infection, inoculation and roguing rates between two plant types affect epidemic potential, depending on the value of *R*_*P*_.

**Fig. 2:**
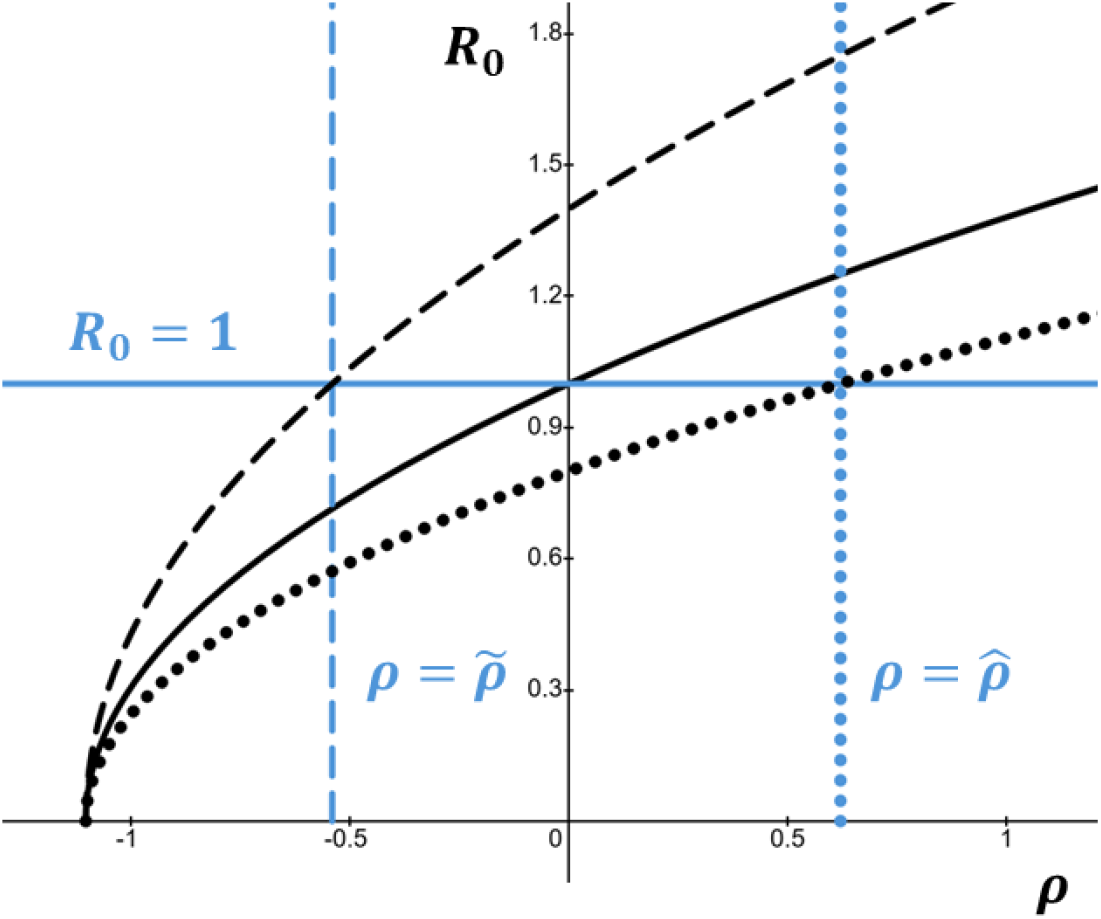
*R*_0_ as a function of *ρ* (black lines) for *R*_*P*_ = 1 (solid), *R*_*P*_ = 0.8 *<* 1 (dotted), and *R*_*P*_ = 1.4 *>* 1 (dashed), with *δ* = 0.9. The blue lines are respectively *R*_0_ = 1 (solid), 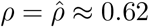 (dotted) and 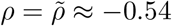 (dashed).

From Fig. 2 we can deduce the following:

– If *R*_*P*_ *<* 1, then the baseline potential for virus spread is low (when *ρ* = 0). Even if the intercrop is more susceptible or less well managed (*ρ >* 0), the system may still suppress disease spread, so long as *ρ* is not too large. We can see in Fig. 2 that there also exists 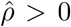 such that *R*_0_ *>* 1 for all 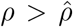. This implies that even modest increases in the susceptibility or mismanagement of the intercrop can tip the system into an epidemic regime. Thus, the most effective strategy here would be to introduce any intercrop with similar or lower susceptibility, but avoid highly susceptible intercrops, which can tip the system into a persistent outbreak regime.
– If *R*_*P*_ *>* 1, then we have a persistent outbreak in our crop only system (when *ρ* = 0). Therefore, introducing a similarly or more susceptible intercrop (*ρ >* 0) will only make this worse. However, if the intercrop is less susceptible or more aggressively managed (*ρ <* 0), it may help reduce *R*_0_, but only if this difference is sufficiently large. We can see in Fig. 2 that there also exists a threshold 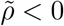 such that *R*_0_ *>* 1 for all 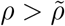. The most effective strategy here would be to only introduce a high proportion of intercrops with substantially lower susceptibility or more aggressive management (*ρ* ≪ 0).

In Fig. 2, for 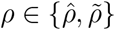, and for a given *δ >* 0 and *R*_*P*_ *>* 0, we can write

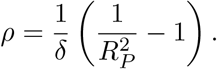

The lines 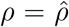 and 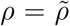 respectively describe the threshold beyond which increasing the relative deviation in viral transmission effort results in persistent disease outbreaks when *R*_*P*_ *<* 1 and *R*_*P*_ *>* 1. The intercropping system tends to be especially sensitive to small changes in parameters near the threshold values 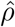 and 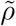, emphasising the importance of careful management to limit disease spread.

### 4.2 Implications for Management Strategies

These results theoretically show that intercropping does not inherently suppress persistent virus outbreaks. The epidemiological outcome depends critically on the intercrop’s infectivity and management relative to the main crop. In some cases, intercropping may inadvertently increase *R*_0_, if the intercrop acts more as a source of viral transmission than as a suppressive barrier. Grauby et al. (2022) observed that intercropping sometimes has little to no effect on reducing Barley Yellow Dwarf Virus transmission in aphids. This empirical finding highlights the importance of intercrop selection and targeted interventions to achieve effective disease suppression. In some cases crops may promote the development of alate (winged) aphids if the crops attracts and retains an aphid pest that harbours a non-persistent virus. This facilitates positive density dependence, inducing aphid dispersal due to intraspecific competition (Carr et al., 2020; Donnelly et al., 2019). These results show that persistent outbreak prevention depends not just on increasing plant diversity, but on the specific management of both crop and intercrop species. Intercrops that transmit infection poorly or are removed quickly suppress *R*_0_, while highly susceptible or poorly managed intercrops can amplify it.

## 5 Persistent Outbreak Risk in Monoculture vs. Intercropped Fields

In this section, to address Q2, we compare monoculture and intercropping systems by examining how the weighted total plant population sizes affect *R*_0_. Specifically, we analyse the relative change in *R*_0_ when introducing intercrops to assess how added host diversity alters epidemic potential. In the Supplementary Material we included time series examples when a persistent outbreak in a monoculture system can be stabilised by introducing an intercrop. We also included the opposite behaviour, when a stable disease-free equilibrium in a monoculture system can be destabilised by introducing an intercrop.

Recall from (4) that 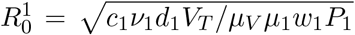 is the basic reproduction number for a monoculture system with *P*_1_ plants. We can calculate the relative change in *R*_0_ when we move from a monoculture to an intercropped system as

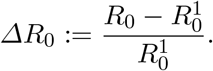

We can write this expression simply in terms of *ρ*_*i*_, *w*_*i*_ and *P*_*i*_ as

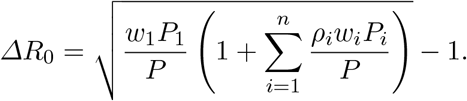

Using this expression, we will now look at the *n* = 2 case, to investigate how changing *w*_1_*P*_1_ and *w*_2_*P*_2_ alters the basic reproduction number in a monoculture and intercropped system, as well as how *ΔR*_0_ changes, for various *ρ* values. This is to show how varying total plant densities and/or susceptible plant attractiveness results in various *R*_0_ regimes. Three disctinct scenarios arise depending on the magnitude and sign of *ρ*, as seen when we look at *ΔR*_0_ = 0 in the (*w*_1_*P*_1_, *w*_2_*P*_2_)-plane (see the Supplementary Material for details).

### 5.1 Intercrops as Barriers to Disease Spread

If *ρ <* 0 the intercrop acts as a barrier to disease spread, transmitting infection less effectively than the crop. The line *ΔR*_0_ = 0 has negative slope in this case and so it never intersects the positive quadrant. In Fig. 3 we see the regimes that emerge when *ρ <* 0. There is a finite region for relatively low enough weighted crop and intercrop densities where persistent outbreaks occur in both monoculture and intercropped fields. Two additional regions are prominent in this scenario. For relatively low enough weighted plant densities, introducing sufficiently high weighted densities of intercrops causes a shift from a persistent outbreak regime under a monoculture to a stable disease-free equilibrium when intercropping, confirming intercropping’s beneficial role. When both *w*_1_*P*_1_ and *w*_2_*P*_2_ are large enough, stability is achieved in both monoculture and intercropped systems, likely because the vector population becomes insufficient to maintain transmission either across a large host community and/or acorss a community where healthy crops are largely preferred.

**Fig. 3:**
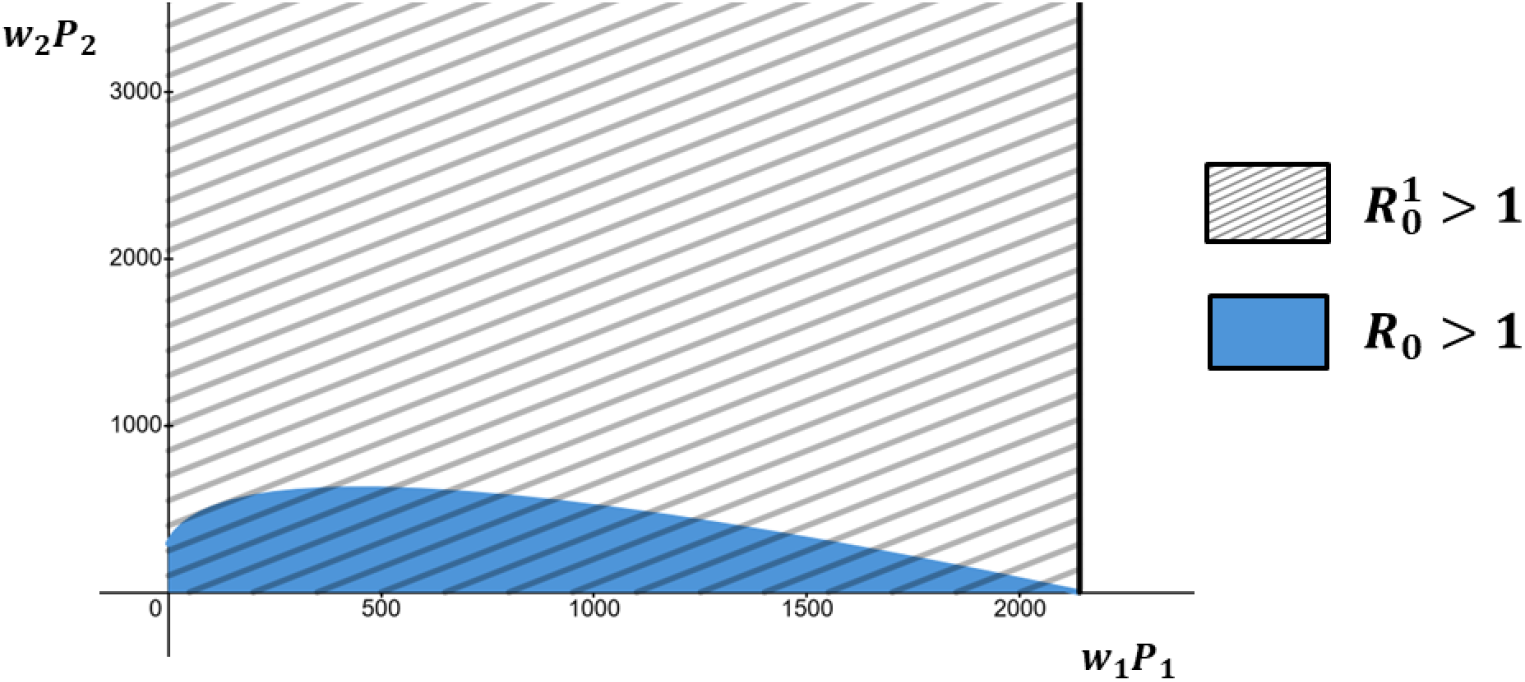
The general behaviour of how the basic reproduction number changes when moving from a monoculture 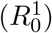 to an intercropped system (*R*_0_) when varying *w*_1_*P*_1_ and *w*_2_*P*_2_, where *ρ <* 0. The blue region is where *R*_0_ *>* 1 (outbreak for intercropped system). In this scenario we have that *ΔR*_0_ *<* 0 for all *w*_1_*P*_1_ and *w*_2_*P*_2_ (intercropping always decreases disease risk). The solid black line is where 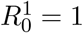. The the left of this line, the shaded grey area, is where 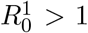 (outbreak in monoculture). To the right of this line is where 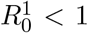. Parameters: *c*_1_ = 2, *c*_2_ = 0.3, *d*_1_ = 6.2, *d*_2_ = 7, *ν*_1_ = 1.7, *ν*_2_ = 1, *µ*_1_ = 1.9, *µ*_2_ = 1.4, *V*_*T*_ = 270 and *µ*_*V*_ = 1.4.

### 5.2 Intercrops as Facilitators of Disease Spread

If *ρ >* 0, the intercrop facilitates disease spread more than the crop, possibly by promoting vector movement or increasing contact rates. For *ρ* ∈ [0, 1] we observe similar behaviour to when *ρ <* 0. The line *ΔR*_0_ = 0 has negative slope in this case and so it never intersects the positive quadrant. In Fig. 4 we can see the regimes that emerge when *ρ* ∈ (0, 1]. There is a finite region for relatively low crop and intercrop densities where persistent outbreaks occur in both monoculture and intercropped fields. As we increase *w*_1_*P*_1_ we generally transition from an outbreak regime in the monoculture to a stable disease-free equilibrium regime in the monoculture system. For a large range of *w*_2_*P*_2_ values we get that the intercropped system has a stable disease-free equilibrium. This however depends on how large the total plant densities are and how attractive susceptible intercrops are.

**Fig. 4:**
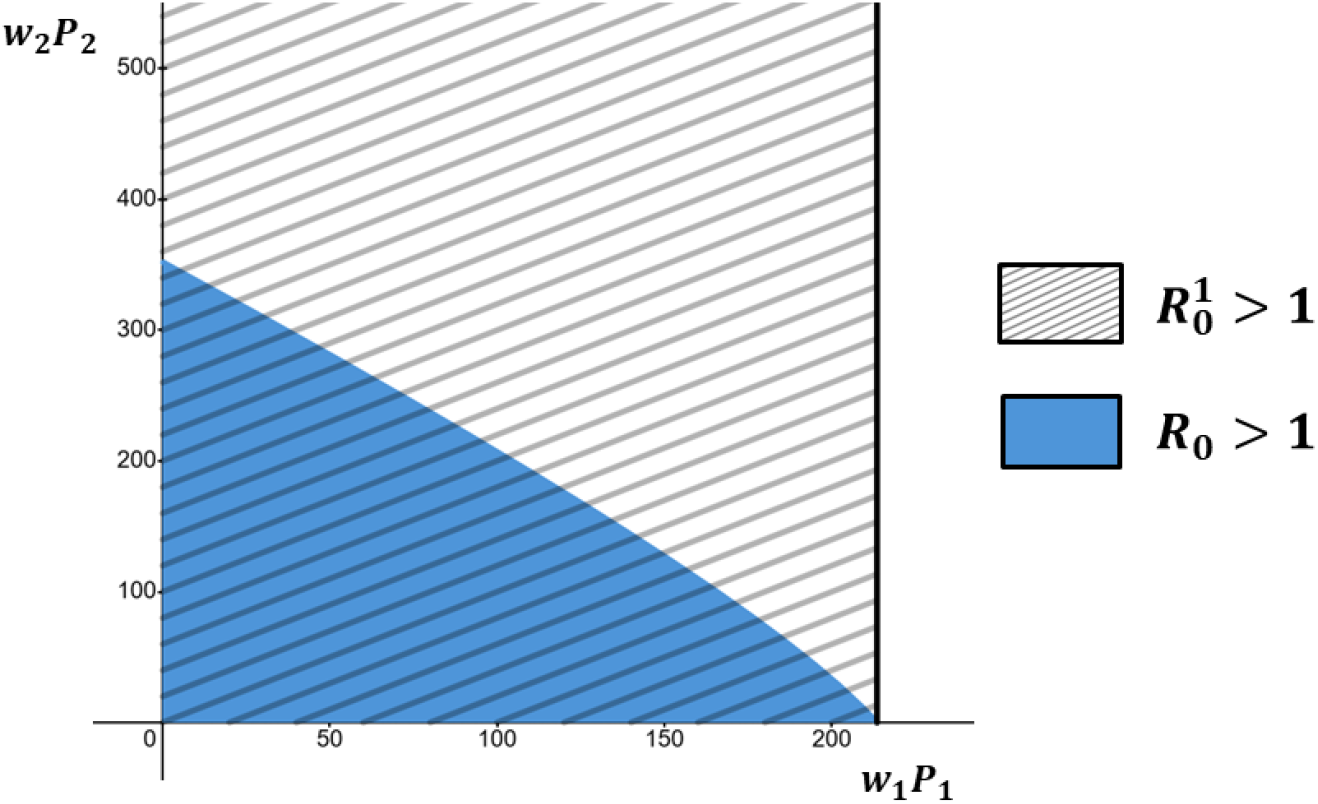
The general behaviour of how the basic reproduction number changes when moving from a monoculture 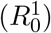 to an intercropped system (*R*_0_) when varying *w*_1_*P*_1_ and *w*_2_*P*_2_, where *ρ* ∈ [0, 1]. The blue region is where *R*_0_ *>* 1 (outbreak for intercropped system). In this scenario we have that *ΔR*_0_ *<* 0 for all *w*_1_*P*_1_ and *w*_2_*P*_2_ (intercropping always decreases disease risk). The solid black line is where 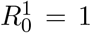. The left of this line, the shaded grey area, is where 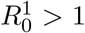 (outbreak in monoculture). To the right of this line is where 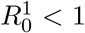. Parameters: *c*_1_ = 2, *c*_2_ = 0.5, *ν*_1_ = 0.25, *ν*_2_ = 1.2, *d*_1_ = 6.2, *d*_2_ = 7, *µ*_1_ = 2.8, *µ*_2_ = 2.3, *V*_*T*_ = 270 and *µ*_*V*_ = 1.4.

For *ρ >* 1, we get that the line *ΔR*_0_ = 0 has positive slope and so intersects the interior of the positive quadrant. The line *ΔR*_0_ = 0 is the yellow dashed line in Fig. 5. We can observe that when *w*_1_*P*_1_ is sufficiently low, persistent outbreaks occur in the monoculture and persist at low to intermediate *w*_2_*P*_2_ values, indicating limited disease suppression by the intercrop (see Fig. 5). As *w*_2_*P*_2_ increases, the basic reproduction number initially decreases then rises, but both monculture and intercropped systems have a disease-free equilibrium that eventually stabilises with persistent outbreaks unlikely to occur. At high *w*_2_*P*_2_ values, the system shifts from a persistent outbreak regime in the monoculture system to a stable disease-free equilibrium regime in the intercropped system, suggesting that a high total density of intercrops and/or a highly attractive healthy sub-population of intercrops can suppress disease spread despite their facilitation of transmission. Interestingly, we can see in Fig. 5 an intermediate region near the line 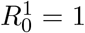, where adding intercrops and/or increasing the attractiveness of susceptibles may cause persistent outbreaks, even though the monoculture had a stable disease-free equilibrium, revealing complex crop-intercrop interactions. Finally, at sufficiently large *w*_1_*P*_1_ and *w*_2_*P*_2_, the disease-free equilibrium stabilises for both systems.

**Fig. 5:**
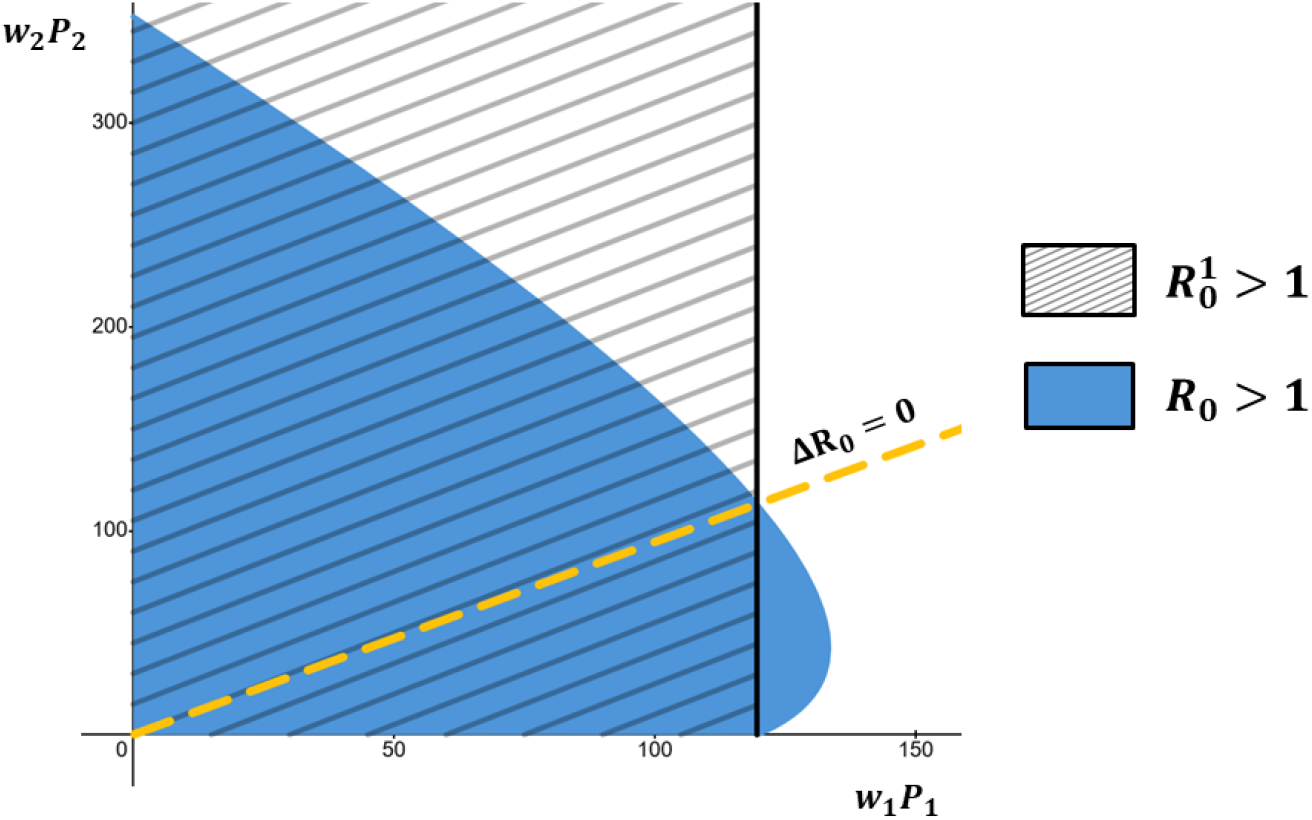
The general behaviour of how the basic reproduction number changes when moving from a monoculture 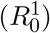 to an intercropped system (*R*_0_) when varying *w*_1_*P*_1_ and *w*_2_*P*_2_, where *ρ >* 1. The blue region is where *R*_0_ *>* 1 (persistent outbreak for intercropped system). The dashed gold line is where *ΔR*_0_ = 0, (where 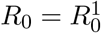). Below and above this line is respectively where *ΔR*_0_ *>* 0 (intercropping increases disease risk) and *ΔR*_0_ *<* 0 (intercropping decreases disease risk). The solid black line is where 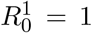. The hatched area, to the left of this line, is where 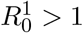 (persistent outbreak in monoculture). To the right of this line is where 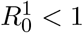. Parameters: *c*_1_ = 0.25, *c*_2_ = 1.2, *ν*_1_ = 2, *ν*_2_ = 0.5, *d*_1_ = 6.2, *d*_2_ = 7, *µ*_1_ = 5, *µ*_2_ = 2.3, *V*_*T*_ = 270 and *µ*_*V*_ = 1.4.

### 5.3 Implications for Management Strategies

The results in this section broadly align with the ecological concept of the concentration effect, where decreasing host density or diversity can increase disease risk by concentrating the virus within a small host community or concentrating the impact of highly competent hosts. In addition to these direct effects on transmission, intercropping can influence pathogen dynamics through so-called trait-mediated indirect effects, changes in vector behavior that modify transmission rates without altering host densities (van Veen et al., 2005). Such indirect effects can promote stable coexistence in host-vector communities by balancing interactions, and in our context, they may inhibit disease spread, due to the possibly higher attractiveness of the healthier susceptible community. Our model captures trait-mediated indirect effects by reflecting how weighted intercrop densities alters transmission rates via *w*_*i*_*P*_*i*_. We have shown that the epidemiological outcome depends not only on host transmission competence but also on their relative abundances and susceptible attractiveness properties. When the intercrop is a poor transmitter, increasing its density can lower *R*_0_, stabilising the disease-free equilibrium. This can also be achieved through increasing the attractiveness of the susceptible sub-populations. However, if the intercrop supports transmission, even modest increases in its population size or susceptible attractiveness can have the potential to raise the basic reproduction above the epidemic threshold. This in turn also depends on the interaction between how large the relative deviation in viral transmission effort is and the size of the total weighted plant densities/abundances.

It is well-known that the basic reproduction number can be generally interpreted as the expected number of secondary infections caused by a single infected individual in a fully susceptible population (Van den Driessche, 2017). We can then see from the above results that not only does the value of *R*_0_ and 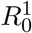 matter, but also the value of *ΔR*_0_, as this captures how intercropping can change the number of secondary infections that occur on average from an individual plant or vector. When *ρ <* 0 or *ρ* ∈ [0, 1] we can see that *ΔR*_0_ *<* 0, i.e. intercropping always decreases disease risk, for any chosen *w*_1_*P*_1_ and *w*_2_*P*_2_. Thus, the best strategy here is introducing intercrops that are barriers to disease spread. One can also introduce intercrops that are facilitators to disease spread, but they must not differ substantionally to the main crop of interest in terms of their transmission characteristics. We can also see that, even in the regime where persistent outbreaks occur in both monoculture and intercropped systems in Fig. 3, *ΔR*_0_ *<* 0 indicates that intercropping lowers the basic reproduction number, thus reducing the number of expected secondary infections by individual hosts or vectors. Lowering *R*_0_ via intercropping is a useful control strategy for managers and farmers, as it can reduce crop yield losses due to infection. This is an important practical consideration, as in real-world systems *R*_0_ may only be lowered but still remain above unity. For example, this kind of behaviour has been observed in relation to cassava mosaic virus, where intercropping cassava with cowpea or maize can reduce disease incidence compared with cassava monocultures, even though the disease still persists within each system (Fondong et al., 2002).

## 6 Transient Outbreak Potential Despite Possible Disease Extinction

From an applied perspective, population densities are observed over finite time intervals, making short-term dynamics as important as long-term qualitative behaviour determined by *R*_0_ (Li & Zou, 2024). Although *R*_0_ *<* 1 ensures eventual decline of infected densities for small enough initial densities, it does not rule out transient amplifications, temporary increases in infection following small perturbations or disturbances. For example, an initial surge in infections can still cause significant crop yield losses within a single growing season. To characterise this behavior, and to provide an answer to Q3, we will analyse such a system’s reactivity, which quantifies when there is the high possibility of having an initial growth rate of our system following small enough perturbations from the disease-free equilibrium (Arnoldi et al., 2016; Hosack et al., 2008; Trevisin et al., 2022). Reactivity is especially relevant in epidemiology, as if transient outbreaks are realised in reactive systems, these can cause substantial harm (O’Regan et al., 2020).

Positive reactivity, *R*_*J*_ *>* 0, indicates that there exists at least one initial condition that results in transient outbreaks (Hosack et al., 2008). The opposite of positive reactivity, so when *R*_*J*_ *<* 0, is called initial resilience, where trajectories cannot exhibit any transient growth, but strictly decrease toward the extinction equilibrium. If one knows their system is reactive, whether a transient outbreak occurs or not depends on the initial conditions chosen. As outlined by Mari et al. (2017) one will observe transient outbreaks if their system has positive reactivity and their initial conditions are contained within the so-called *g*-reactivity basin, which can be constructed using the Jacobian of your system (see the Supplementary Material for more details).

### 6.1 The Threshold Index for Epidemicity

We now look at characterising transient outbreak potential by deriving the threshold index for epidemicity ℰ_0_. The quantity ℰ_0_ is called a threshold index for epidemicity if ℰ_0_ *>* 1 ⇒ *R*_*J*_ *>* 0 and ℰ_0_ *<* 1 ⇒ *R*_*J*_ *<* 0. Therefore ℰ_0_ acts as an indicator, telling us when transient outbreaks are possible, or when trajectories strictly decrease toward the extinction equilibrium. Hosack et al. (2008) showed that the threshold index for epidemicity can be computed as

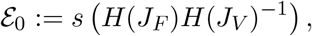

where *H*(*A*) is the Hermitian part of a matrix *A*. The matrix *H*(*J*_*F*_)*H*(*J*_*V*_)^−1^ is called the maximum next-generation matrix, where *J*_*F*_ and *J*_*V*_ are the same as in (3). ℰ_0_ is interpreted as the maximum number of new infections produced by an infective individual at the disease-free equilibrium (Hosack et al., 2008).

For our model, we can compute that (see Supplementary Material for derivation)

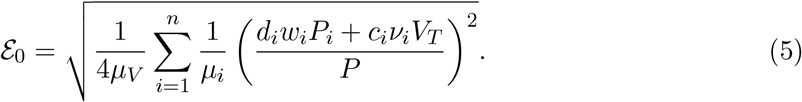

For *n* = 2, observing the behaviour of ℰ_0_ in the (*c*_1_, *c*_2_)-, (*d*_1_, *d*_2_)-, (*ν*_1_, *ν*_2_)-, or (*w*_1_, *w*_2_)-plane reveals a pattern similar to how *α*_*i*_ affects *R*_0_ (see Fig. 1). For example, larger acquisition or inoculation rates induce positive reactivity, while smaller values lead to initial resilience, showing that, in general, more efficient virus transmission by the vector or plant increases transient outbreak potential.

### 6.2 Total Weighted Plant Densities, Reactivity and Initial Resilience

When we observe ℰ_0_ in the (*w*_1_*P*_1_, *w*_2_*P*_2_)-plane, distinct initial resilience, reactive and persistent outbreak regions emerge, which depends on the parameters of our model (see Fig. 6, Fig. 7 and the Supplementary Material). The behaviour of these regions can be inferred using conic sections (see the Supplementary Material for derivations). There are two main scenarios that emerge:

**Fig. 6:**
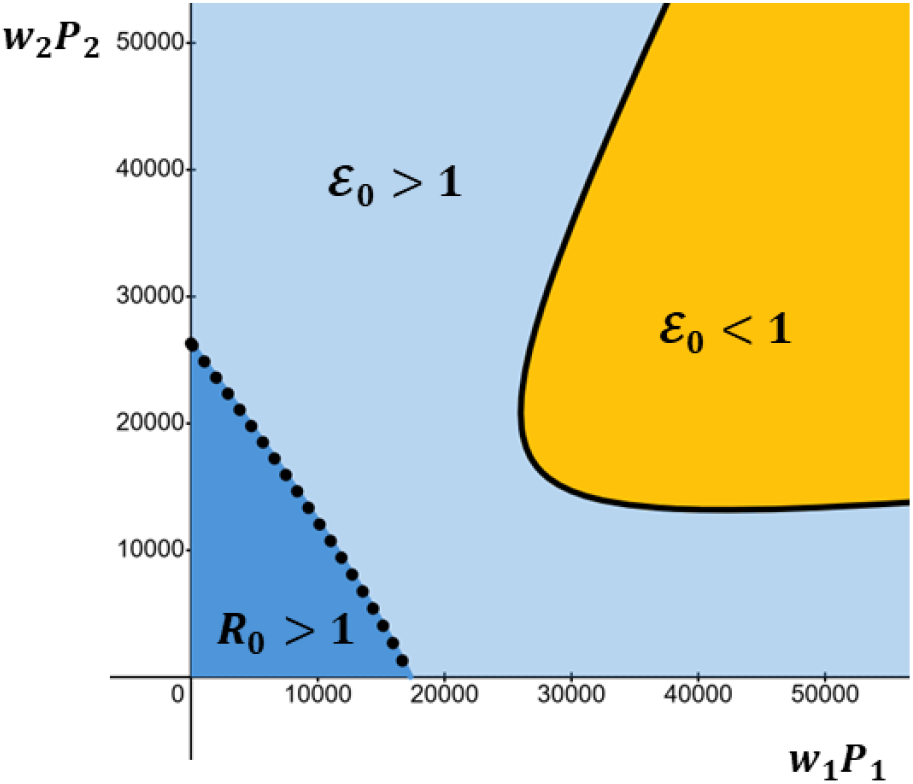
Illustrative example of the (*w*_1_*P*_1_, *w*_2_*P*_2_)-plane where we can observe a reactive region (light blue), an initially resilient region that does not intersect either axis (yellow), and a region where persistent outbreaks occur (dark blue). The solid black curve is ℰ_0_ = 1 and the dotted black curve is *R*_0_ = 1. Parameters: *c*_1_ = 4, *c*_2_ = 3.6, *ν*_1_ = 2, *ν*_2_ = 1, *d*_1_ = 11, *d*_2_ = 5.2, *µ*_1_ = 10, *µ*_2_ = 1.4, *V*_*T*_ = 4920 and *µ*_*V*_ = 2.5.

**Fig. 7:**
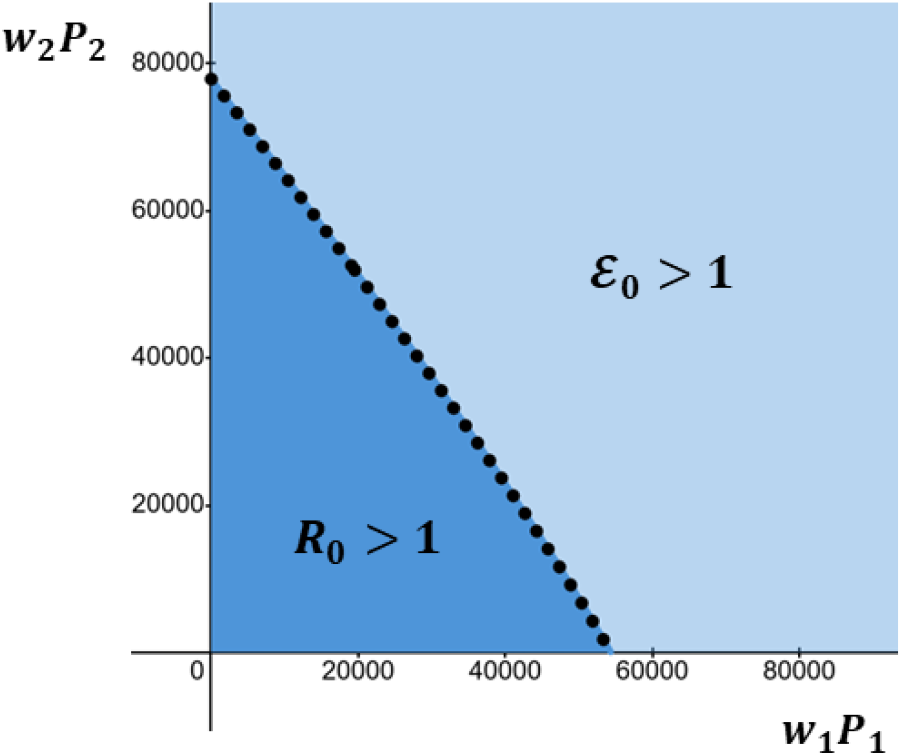
Illustrative example of the (*w*_1_*P*_1_, *w*_2_*P*_2_)-plane where there is only a reactive region (light blue) and a region where persistent outbreaks occur (dark blue). The dotted black curve is *R*_0_ = 1. Parameters: *c*_1_ = 4, *c*_2_ = 3.6, *ν*_1_ = 2, *ν*_2_ = 1, *d*_1_ = 7.6, *d*_2_ = 7.7, *µ*_1_ = 2.2, *µ*_2_ = 0.7, *V*_*T*_ = 4920 and *µ*_*V*_ = 2.5.

1. There exists at least one and at most two reactive regions, where transient outbreaks are possible. This occurs precisely when

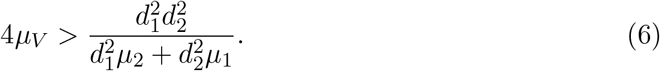

When (6) holds a reactive region lies between a persistent outbreak region and a region of initial resilience. There are several sub-cases that can occur, which depend on whether the curve ℰ_0_ = 1 intersects the positive *w*_*i*_*P*_*i*_-axis. This occurs precisely when

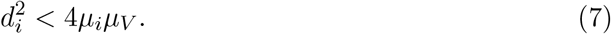

An interesting sub-case can be seen in Fig. 6, when ℰ_0_ = 1 does not intersect either the *w*_1_*P*_1_- and *w*_2_*P*_2_-axis. This occurs precisely when (7) does not hold. In this case, intercropping is essential to allow initial resilience to emerge, as when the crop and intercrop are in a monoculture system only a persistent outbreak region and reactive region exist for all choices of *w*_1_*P*_1_ and *w*_2_*P*_2_. More sub-cases are detailed in the Supplementary Material, for example when ℰ_0_ = 1 intersects the *w*_*i*_*P*_*i*_-axis. When this occurs, this means that initial resilience can emerge for the corresponding plant host when in a monoculture, if the total weighted plant *i* density is sufficiently large. In general, regardless of the sub-case, when a region of initial resilience exists, we can observe that at sufficiently low total plant densities, be it the intercrop or crop, persistent outbreaks occur. This is perhaps due to concentration effects, where decreasing host densities concentrates the virus in the host population (Wonham et al., 2006). Moderate to high transmission rates can shrink the initial resilience region, in the sense that larger total densities are needed for initial resilience to emerge. Low host transmission rates (low *d*_2_, for example) can substantially reduce transient outbreak potential, especially at higher *w*_*i*_*P*_*i*_ values. Thus, host plants with low transmission efficiency can enhance initial resilience by having the potential to prevent early epidemic spikes. Increasing infected plant or infective vector loss rates (*µ*_*i*_ and *µ*_*V*_) can decrease ℰ_0_ and ultimately narrow the reactive region, as can be deduced from (7). This implies that initial resilience can occur over a wider range of both total weighted plant densities and susceptible attractiveness values.
2. There only exists a reactive region and a persistent outbreak region (Fig. 7). This occurs whenever (6) does not hold. In this case, ℰ_0_ = 1 does not intersect the positive quadrant where *w*_1_*P*_1_ and *w*_2_*P*_2_ are nonnegative (see the Supplementary Material for more details). In this scenario, at sufficiently high *w*_1_*P*_1_ and *w*_2_*P*_2_ one enters an unbounded reactive region and for lower *w*_1_*P*_1_ and *w*_2_*P*_2_ persistent outbreaks emerge. This scenario means that in both the monoculture and intercropping systems, there is potential for no outbreak to occur, be it transient or persistent, but as demonstrated above, this further depends on whether or not the initial conidtions are in the *g*-reactivity basin. In this scenario there are more weighted mixture sizes that allow for the possibility for outbreaks, as there is no region of initial resilience.

We have thus demonstrated a clear dichotomous pattern, between when a region of initial resilience emerges or not, when varying *w*_1_*P*_1_ and *w*_2_*P*_2_, which relies solely on (6) holding or not. In order for a region of initial resilience to emerge we must have that the vector loss rate is sufficiently large relative to the rate of plant inoculation. Our results are quite robust, as choosing certain numerical parameter values for model simulation will just alter the specific shape and scale of these regions. We conjecture that the above results generalise to *n* ≥ 2 (see the Supplementary Material for more details).

### 6.3 Implications for Management Strategies

This analysis shows that intercropping can influence not only disease persistence but also transient outbreak risk. High-density intercropping and/or highly attractive susceptible plants, with low transmission efficiency, appears to be most favourable for suppressing both persistence and transient outbreak risk. Conversely, introducing intercrops with high infectivity may backfire, enabling the possibility of damaging transient outbreaks even when in the long-term one observes disease extinction. The latter scenario may not be from deliberate planting decisions but could instead arise from unintentional factors, such as the invasion of a novel viral strain or shifts in vector behavior due to environmental conditions.

While *R*_0_ remains an important quantity to characterise, it does not capture fully the risks posed by transient outbreak potential in both monoculture and intercropping systems. The presence or absence of initial resilience reflects how robust each system is to small introductions of infective densities and plays an important role in determining which and how plants should be intercropped. In the Supplementary Material we further discuss two biologically motivated examples when studying the behaviour of ℰ_0_, one in relation to the simple case when both crop and intercrop have identical transmission and removal parameters, and one in relation to varietal mixtures. The sensitivity of ℰ_0_ to changes in parameter values can be seen from the several scenarios that emerge when (6) holds. This inequality might hold and so initial resilience emerges, if, for example, the vector population loses the virus faster than it can transmit it to either plant, e.g. for non-persistent viruses (Falla & Cunniffe, 2024); if the crop and/or intercrop have low susceptibility, e.g. resistant/non-resistant varieties (McAlvay et al., 2022); or if infected plants are removed quickly, e.g. using sentinel plants (Lovell-Read et al., 2023).

We found that in certain contexts, intercropping may be essential to allow initial resilience to occur, precisely when (7) does not hold. That is, if two crops are in a monoculture system where the plant inoculation rate is sufficiently high, we may only observe persistent and transient outbreaks (depending on the initial condition chosen). But when they are inter-cropped we can find mixture compositions that yield initial resilience, as illustrated in Fig. 6. A homogenous population within a monoculture could facilitate the rapid amplification of virus spread, whereas intercropping with mixed densities has the potential to introduce sufficient heterogeneity that may disrupt host continuity and ultimately allow initial resilience to emerge.

## 7 Discussion

Our theoretical analysis provides key insights into vector-borne plant virus transmission in intercropping systems, showing how pathogen evolution in heterogeneous host environments can alter the potential for disease outbreaks. We derived and interpreted both the basic reproduction number, *R*_0_ and the epidemicity index, E_0_, which together offer a nuanced understanding of how different intercrop choices alter disease dynamics. Importantly, our results highlight that high-density, low-transmission intercrops can suppress persistent outbreaks and limit the likelihood of transient outbreaks. This complements empirical studies showing intercropping often reduces disease incidence by disrupting vector movement or diluting susceptible hosts (Grauby et al., 2022; Hooks & Fereres, 2006; Roudine et al., 2025; Tous-Fandos et al., 2025).

Our results also imply that in some systems, especially with novel or poorly characterised viruses, intercropping can backfire. For example, if a virus infects multiple hosts or its vector is highly mobile, mixed plantings can increase transmission opportunities and maintain infective vector populations, thus amplifying rather than suppressing epidemics. Overall, these findings underscore that the success of intercropping depends not merely on adding diversity, but on careful selection of intercrop species based on their ecological and epidemiological roles. Intercropping designs informed by mechanistic models such as ours have the potential to more effectively enhance sustainability and disease control. A particularly interesting result is the example in Fig. 6, where larger mixture sizes or higher susceptible attractiveness of the two hosts can have greater potential to suppress transient outbreaks that would otherwise be highly likely in isolated monocultures. We have not found real world examples where this effect has been reported, but it would be interesting to see if this result can be empirically validated, either with field or laboratory experiments. Despite this, it is a useful theoretical insight that can help guide empirical studies by suggesting specific host mixtures to test for potential transient outbreak suppression in real-world settings.

Our study provide several opportunities for future research. Mathematically, our analysis focused on local behavior near the disease-free equilibrium, where *R*_0_ *<* 1 leads to extinction for small enough initial infected densities. As mentioned previously, in all observed simulation scenarios we conducted (some of which are provided in the Supplementary Material), *R*_0_ *<* 1 resulted in disease extinction for all chosen positive initial conditions. Proving this mathematically, and also providing sufficient conditions for global stability for equilibria of (1), remain as open problems. One key simplification of our model is the assumption of a constant total plant and vector population sizes. In both natural and agricultural systems, plant populations are dynamic, subject to growth, mortality, and competitive interactions. Introducing explicit population dynamics and plant competition could reveal feedbacks between ecological and epidemiological processes that affect virus persistence. We also assume only horizontal transmission of viruses. To extend our framework it would be interesting to incorporate vertical virus transmission (Pagán et al., 2014).

In our framework, infected or susceptible plants may be more attractive for certain vector species. We incorporated this into the model as conditional vector preferences. We looked at how varying the weighted plant densities, *w*_*i*_*P*_*i*_, affected disease dynamics. There are many cases to consider for different susceptible/infected attractiveness plants. It would be interesting for future studies to look at vector preference scenarios in the context of intercropping. This would follow in a similar vein to the work of Falla and Cunniffe (2025), who studied a detailed model of virus control within various companion planting scenarios. Note that McElhany et al. (1995) highlighted the relationship between vector behavior and disease spread can also be density-dependent. For example, for Barley Yellow Dwarf Virus, when diseased plants are rare, preference for infected plants promotes transmission, whereas at high disease prevalence, preference for healthy plants has a stronger effect. Moreover, vector preferences are known to differ substantially between persistent and non-persistent viruses (Cunniffe et al., 2021). Exploring these various aspects within our framework would further elucidate how conditional vector preferences affect outbreak dynamics.

Spatial scale and the arrangement of crops play a crucial role in shaping disease dynamics, as vector movement and pathogen transmission often depend on the spatial configuration of host plants (Allen-Perkins & Estrada, 2019; Hamelin et al., 2023; McLeish et al., 2017; Rother et al., 2025). Different cropping patterns, such as mixed or row intercropping, create heterogeneous landscapes that may either suppress or enhance virus spread. In practice, virus transmission is also affected by a range of environmental and management factors, such as temperature-dependent vector activity, and the timing and efficacy of insecticide applications, for example (Duffy et al., 2017; Liang et al., 2012). Incorporating such aspects would allow the model to generate more context-specific insights and could inform precision agriculture strategies. Further realism could also be achieved by parameterising the model for specific crop-virus-vector systems. Another critical next step is to link this theoretical framework with empirical data. Calibrating the model using field or experimental observations would allow for parameter estimation and validation of predictions (Kendall et al., 1992). In our framework pathogen transmission is assumed to occur instantaneously. It would be interesting to extend our intercropping framework to incorporate time delays or additional infection classes (such as exposed or recovered compartments). Such extensions could capture greater biological realism, including latent infectious periods between vectors and hosts (Brauer et al., 2019).

An important aspect of using mathematical models is their real-world applicability. The model we propose not only allows for the study of intercropping different hosts. It can also be used to assess how mixing resistant and susceptible crops within a varietal mixture of the same species influences pathogen transmission (Newton & Skelsey, 2023). In intercropping systems, one may intercrop plants to deter or attract vectors associated with a main crop of interest. Additionally, our framework could be used to model the intercropping of two or more economically important crops that may experience outbreaks when grown as monocultures but maintain a stable disease-free equilibrium when intercropped. In such cases, both crops are expected to achieve higher yields compared to when they are grown in isolated fields (Fondong et al., 2002).

## Acknowledgements

We would like to thank Louise McNamara, Stephen Byrne, and Maximilian Schughart for their insightful conversations at the early stages of this project.

