## Supplementary Material for "Disentangling the Effects of Intercropping on Vector-Borne Plant Virus Dynamics"

In this Supplementary Material we derive the main expressions found in the original manuscript and give the necessary theoretical background for understanding our modelling framework. We also provide additional plots that give more insight into the dynamical behaviour our model can capture. All numerical simulations were conducted in R version 4.5.1 (R Core Team, 2025). The code used can be provided upon request.

For $n \in \{1, 2, ...\}$, we will first define the positive and negaitive orthants (quadrants if $n = 2$) respectively as

$$\mathbb{R}^n_+ := \{x \in \mathbb{R}^n : x_i \geq 0\},$$
$$\mathbb{R}^n_- := \{x \in \mathbb{R}^n : x_i \leq 0\}.$$

The interior of these sets are given by

$$\text{Int}(\mathbb{R}^n_+) := \{x \in \mathbb{R}^n_+ : x_i > 0\},$$
$$\text{Int}(\mathbb{R}^n_-) := \{x \in \mathbb{R}^n_- : x_i < 0\}.$$

### 1 The Multi-Host Model

The system we consider in the original manuscript comprises of $n \geq 2$ distinct crop species. An individual crop can be either infected or susceptible (i.e. not infected), where $P_{k,I}$ and $P_{k,S}$ are respectively the number of infected and susceptible individuals for crop $k$). The vector population is likewise divided into susceptible ($V_S$) and infected ($V_I$) compartments. For $k \in \{1, ..., n\}$, the coupled model is given by

$$\begin{aligned}
\frac{dP_{k,S}}{dt} &= \mu_k P_{k,I} - \frac{d_k w_k V_I P_{k,S}}{T(P_S, P_I)}, \\
\frac{dP_{k,I}}{dt} &= \frac{d_k w_k V_I P_{k,S}}{T(P_S, P_I)} - \mu_k P_{k,I}, \\
\frac{dV_S}{dt} &= \mu_V V_I - \frac{V_S \sum_{i=1}^n c_i \nu_i P_{i,I}}{T(P_S, P_I)}, \\
\frac{dV_I}{dt} &= \frac{V_S \sum_{i=1}^n c_i \nu_i P_{i,I}}{T(P_S, P_I)} - \mu_V V_I,
\end{aligned} \tag{1}$$

with the initial condition $X(0) \in \mathbb{R}_+$, for $X \in \{P_S, P_I, V_S, V_I\}$, where $P_S := \{P_{1,S}, ..., P_{n,S}\}$, $P_I := \{P_{1,I}, ..., P_{n,I}\}$, and

$$T(P_S, P_I) := \sum_{i=1}^n (w_i P_{i,S} + \nu_i P_{i,I}).$$

We can see that

$$\frac{dP_{k,S}}{dt} + \frac{dP_{k,I}}{dt} = 0$$
$$\frac{dV_S}{dt} + \frac{dV_I}{dt} = 0.$$

This implies that $P_{k,S} + P_{k,I}$ and $V_S + V_I$ are constant over time. Let $P_{k,S} + P_{k,I} = P_k$ and $V_S + V_I = V_T$ for some constants $P_k \geq 0, V_T > 0$.

### 2 Positivity of Solutions

A fundamental requirement for models to be ecologically meaningful is that solutions remain nonnegative over time, provided that the initial conditions are nonnegative. We can show this using the following standard result.

**Theorem 1.** *(Haddad et al., 2010) Let $f : \mathbb{R}^n \to \mathbb{R}^n$ be a smooth function. Then, the following are equivalent:*

*(a) The nonnegative orthant $\mathbb{R}^n_+$ is forward invariant under the flow of the system*

$$\frac{dx}{dt} = f(x), \quad x(0) \in \mathbb{R}^n_+.$$

*(b) For all $x \in \mathbb{R}^n_+$ and any index $k$ such that $x_k = 0$, we have $f_k(x) \geq 0$.*

Examining the RHS of system (1), we can see that each $dX/dt$ satisfies

$$X = 0 \implies \frac{dX}{dt} \geq 0,$$

where $X \in \{P_S, P_I, V_S, V_I\}$. Hence, by Theorem 1, the nonnegative orthant is invariant under solutions of (1) and so we can see that, for all $t \geq 0$,

$$P_{k,S}(0), P_{k,I}(0), V_S(0), V_I(0) \geq 0 \implies P_{k,S}(t), P_{k,I}(t), V_S(t), V_I(t) \geq 0.$$

### 3 The Basic Reproduction Number

The following is called the next-generation approach for deriving the basic reproduction number, as found in Van den Driessche and Watmough (2002). Consider the system

$$\frac{dx_i}{dt} = F_i(x) - V_i(x), \tag{2}$$

where $x_i(0) \in \mathbb{R}_+$, $F_i : \mathbb{R}^n_+ \to \mathbb{R}_+$ represents the rate at which new infections appear in compartment $i$, and $V_i : \mathbb{R}^n_+ \to \mathbb{R}_+$ represents the rate at which individuals leave compartment $i$. Assume that both $F_i$ and $V_i$ are smooth. Suppose that $m < n$ compartments correspond to infected compartments and that the system admits a disease-free equilibrium $x^*$. Further, assume that the equations for $x_1, \ldots, x_m$ can be linearised at $x^*$ independently of the remaining compartments, and that $F_i = 0$ for all $i$ with $m + 1 \leq i \leq n$, which means no new infections occur in the uninfected compartments. Let $J_F$ and $J_V$ denote the Jacobian matrices of $F$ and $V$ evaluated at the disease-free equilibrium $x^*$. The basic reproduction number is then defined as

$$R_0 := s(J_F J_V^{-1}),$$

where $s(A)$ is the spectral radius of matrix $A$. We can now state the next well-known result.

**Theorem 2.** *(Van den Driessche & Watmough, 2002) If $x^*$ is a disease-free equilibrium of the system above, then $x^*$ is locally asymptotically stable if $R_0 < 1$, and unstable if $R_0 > 1$.*

To derive the basic reproduction number for our model, we follow the above next-generation matrix approach. We can see that

$$(P_S, P_I, V_S, V_I)^T = (P^*, 0, V_T, 0)^T$$

is the so-called disease-free equilibrium of (1), the fixed point where there are no infected compartments present, where $P^* = \{P_1, ..., P_n\}$. Since the total populations are constant, we can reduce our system (1) to

$$\begin{aligned} \frac{dP_{k,I}}{dt} &= \frac{d_k w_i V_I (P_k - P_{k,I})}{T(P^* - P_I, P_I)} - \mu_k P_{k,I}, \\ \frac{dV_I}{dt} &= (V_T - V_I)\left(\frac{\sum_{i=n}^{n} \nu_i c_i P_{i,I}}{T(P^* - P_I, P_I)}\right) - \mu_V V_I. \end{aligned} \tag{3}$$

The dynamics of (3) must stay within realistic bounds, i.e. the infected compartments must not exceed their respective total population sizes. Since each crop and vector population is constant, we thus have that the nonempty set

$$\mathcal{D} := \left\{(P_I, V_I)^T \in \mathbb{R}^{n+1} : 0 \le P_{k,I} \le P_k, 0 \le V_I \le V_T\right\}$$

is a suitable domain of attraction. That is, solutions of (3) remain within $\mathcal{D}$ for all $t \ge 0$.

Let $V = (\mu_1 P_{1,I}, \mu_2 P_{2,I}, ..., \mu_n P_{n,I}, \mu_V V_I)^T$ and

$$F = \begin{pmatrix} \dfrac{w_1 d_1 V_I (P_1 - P_{1,I})}{T(P^* - P_I, P_I)} \\ \vdots \\ \dfrac{w_n d_n V_I (P_n - P_{n,I})}{T(P^* - P_I, P_I)} \\ (V_T - V_I)\left(\dfrac{\sum_{i=1}^n \nu_i c_i P_{i,I}}{T(P^* - P_I, P_I)}\right) \end{pmatrix}. \tag{4}$$

We can respectively compute the Jacobian of $V$ and $F$ (evaluated at $P_{k,I} = V_I = 0$) as $J_V = \text{diag}(\mu_1, \mu_2, ..., \mu_n, \mu_V)$ and

$$J_F = \begin{pmatrix} 0 & \cdots & 0 & \dfrac{w_1 d_1 P_1}{P} \\ 0 & \cdots & 0 & \dfrac{w_2 d_2 P_2}{P} \\ \vdots & \ddots & \vdots & \vdots \\ 0 & \cdots & 0 & \dfrac{w_n d_n P_n}{P} \\ \dfrac{V_T \nu_1 c_1}{P} & \cdots & \dfrac{V_T \nu_n c_n}{P} & 0 \end{pmatrix}, \tag{5}$$

where $P := \sum_{i=1}^n w_i P_i$. The inverse of $J_V$ is simply $J_V^{-1} = \text{diag}(1/\mu_1, 1/\mu_2, ..., 1/\mu_n, 1/\mu_V)$. Therefore we get that the next-generation matrix is given by

$$J_F J_V^{-1} = \begin{pmatrix} 0 & \cdots & 0 & \dfrac{w_1 d_1 P_1}{\mu_V P} \\ 0 & \cdots & 0 & \dfrac{w_2 d_2 P_2}{\mu_V P} \\ \vdots & \ddots & \vdots & \vdots \\ 0 & \cdots & 0 & \dfrac{w_n d_n P_n}{\mu_V P} \\ \dfrac{V_T \nu_1 c_1}{\mu_1 P} & \cdots & \dfrac{V_T \nu_n c_n}{\mu_n P} & 0 \end{pmatrix}. \tag{6}$$

We computed the eigenvalues of this matrix through column expansion when solving $\det(\lambda I - J_F J_V^{-1}) = 0$. This then gives the basic reproduction number

$$R_0 = \sqrt{\frac{V_T}{\mu_V P^2} \sum_{i=1}^n \frac{\nu_i w_i c_i d_i P_i}{\mu_i}}. \tag{7}$$

If we linearise (3) around the disease-free equilibrium, we can study the linearised system

$$\frac{dx(t)}{dt} = Jx(t),\ x(0) \in \mathbb{R}_+^{n+1}, \tag{8}$$

to evaluate the local stability of our system at the disease-free equilibrium, where $J$ is the Jacobian of (3) evaluated at the disease-free equilibrium, and $x$ represents perturbations from the disease-free equilibrium (Hinrichsen & Pritchard, 2005). This disease-free equilibrium is locally asymptotically stable if and only if $\mu(J) < 0$, where $\mu(A)$ is the spectral abscissa of a matrix $A$. However, in many situations, it is more practical to analyse the basic reproduction number $R_0$, instead of determining the local stability properties of an epidemiological system directly via linearisation. The Jacobian matrix of system (3) at the disease-free equilibrium is given by

$$J = \begin{pmatrix} -\mu_1 & 0 & \cdots & 0 & \frac{w_1 d_1 P_1}{P} \\ 0 & -\mu_2 & \cdots & 0 & \frac{w_2 d_2 P_2}{P} \\ \vdots & \vdots & \ddots & \vdots & \vdots \\ 0 & 0 & \cdots & -\mu_n & \frac{w_n d_n P_n}{P} \\ \frac{V_T \nu_1 c_1}{P} & \frac{V_T \nu_2 c_2}{P} & \cdots & \frac{V_T \nu_n c_n}{P} & -\mu_V \end{pmatrix}.$$

Computing the eigenvalues of $J$ is a challenging task. Even in the simplest case of $n = 2$, solving for $\lambda$ using the characteristic polynomial of $J$, $p_J(\lambda)$ is quite intractable and this complexity increases as $n$ gets larger. Therefore, we study $R_0$ as an indicator of local stability, as per Theorem 2.

### 4 $\varDelta R_0$: Monocultures vs Intercropped Systems

We will now show how the three different scenarios observed in Section "*Persistent Outbreak Risk in Monoculture vs. Intercropped Fields*" of the main manuscript arise. These scenarios depend on the magnitude and sign of $\rho$. This can be seen when we look at $\varDelta R_0 = 0$, for $n = 2$, in the $(w_1P_1, w_2P_2)$-plane. This expression is

$$\sqrt{\frac{w_1P_1}{w_1P_1 + w_2P_2}\left(1 + \frac{\rho_1 w_1 P_1}{w_1P_1 + w_2P_2} + \frac{\rho_2 w_2 P_2}{w_1P_1 + w_2P_2}\right)} - 1 = 0.$$

As $\rho_1 := 0$ and $\rho \equiv \rho_2$, we then get that

$$(w_1P_1 + w_2P_2)^2 - w_1P_1(1+\rho)(w_1P_1 + w_2P_2) + \rho(w_1P_1)^2 = 0.$$

We can reduce this to

$$w_2P_2\left(w_1P_1(1-\rho) + w_2P_2\right) = 0.$$

The two solutions of this quadratic equation are $w_2P_2 = 0$ and $w_2P_2 = w_1P_1(\rho - 1)$, which is a straight line passing through the origin in the $(w_1P_1, w_2P_2)$-plane. From this final expression, we can see that for $\rho < 0$ and $\rho \in [0, 1)$, this line has negative slope and $w_2P_2 < 0$, i.e. it can never intersect the interior of the positive quadrant. When $\rho = 1$, this line becomes $w_2P_2 = 0$ and so can never intersect the interior of the positive quadrant. Finally, for $\rho > 1$, this line has positive slope and so $w_2P_2 \geq 0$, i.e. it does intersect the interior of the positive quadrant. These three conclusions when varying $\rho$ can be seen in the main manuscript.

### 5 Simulation Scenarios

From Theorem 2, we know that $R_0 < 1$ implies local asymptotic stability of the disease free equilibrium. This, however, does not always imply asymptotic stability for all initial conditions, i.e. global asymptotic stability. Despite this, we ran hundreds of simulation studies, each with different initial conditions, and these suggest that $R_0 < 1$ implies global extinction of infected

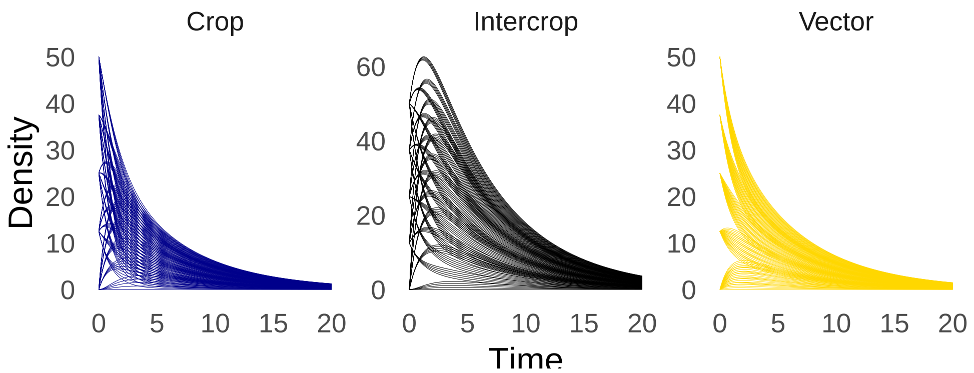


Fig. 1: Example of infected compartment trajectories for a planar version of (3), when $R_0 = 0.73 < 1$, for various initial conditions. Parameters: $d_1 = 1.93$, $d_2 = 1.72$, $P_1 = 10000$, $P_2 = 10000$, $\nu_1 = 1.12$, $\nu_2 = 3.71$, $w_1 = 1$, $w_2 = 2$, $V_T = 500$, $c_1 = 2.93$, $c_2 = 2.2$, $\mu_1 = 0.92$, $\mu_2 = 0.62$, $\mu_V = 0.54$.

compartments for system (3). In Fig. 1 we can see an example of system trajectories when $R_0 < 1$ for various initial conditions, for when $n = 2$. We thus conjecture that $R_0 < 1$ implies global asymptotic stability of the disease-free equilibrium.
In Fig. 2 we can see an example of when a persistent outbreak in a monoculture system can be stabilised by introducing an intercrop.

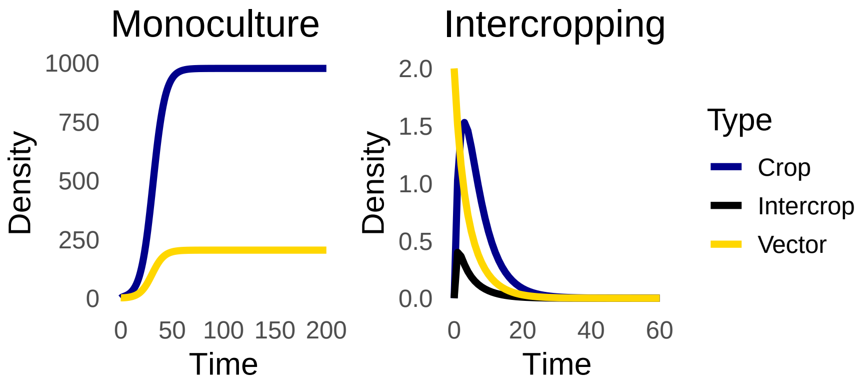


Fig. 2: Stabilisation: Example of infected compartment trajectories for a planar version of (3) for (left) monoculture system and (right) intercropping system. Parameters: $d_1 = 2.6$, $d_2 = 0.75$, $P_1 = 8000$, $P_2 = 10000$, $\nu_1 = 1.5$, $\nu_2 = 1.3$, $w_1 = 1$, $w_2 = 2$, $V_T = 500$, $c_1 = 1.2$, $c_2 = 1.4$, $\mu_1 = 0.45$, $\mu_2 = 1.9$, $\mu_V = 0.3$.

In Fig. 3 we can see an example of when a monoculture system, where infected trajectories go extinct, can be destabilised by introducing an intercrop, inducing a persistent outbreak.

### 6 Metrics for Predicting Transient Dynamics

#### 6.1 Reactivity and the Threshold Index for Epidemicity

Even though $R_0 < 1$ we can still observe short-term outbreaks, despite eventual extinction of infected compartments of (1). Reactivity, denoted $R_J$, measures this short-term growth potential and is defined as

$$R_J := \max_{\|x(0)\|_2=1} \frac{d\|x(t)\|_2}{dt} = \mu(H(J)), \tag{9}$$

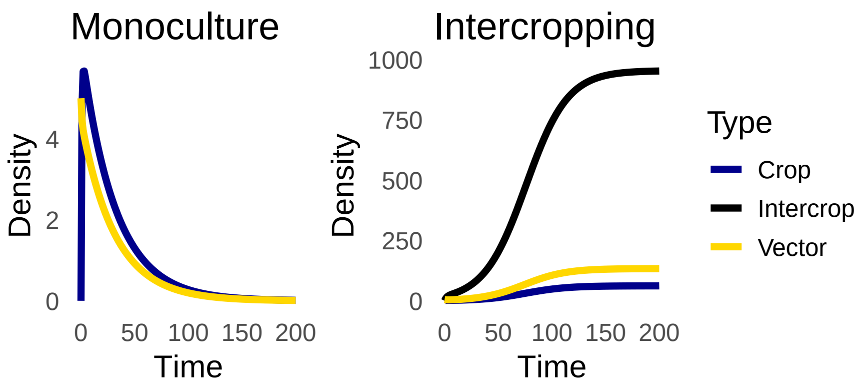


Fig. 3: Destabilisation: Example of infected compartment trajectories for a planar version of (3) for (left) monoculture system and (right) intercropping system. Parameters: $d_1 = 2$, $d_2 = 3.5$, $P_1 = 10000$, $P_2 = 10000$, $\nu_1 = 1.2$, $\nu_2 = 1.5$, $w_1 = 1$, $w_2 = 2$, $V_T = 500$, $c_1 = 2$, $c_2 = 1.4$, $\mu_1 = 1.45$, $\mu_2 = 0.3$, $\mu_V = 0.2$.

where $\|A\|_2$ is the Euclidean norm of $A$ and

$$H(A) := \frac{A + A^T}{2}$$

is the Hermitian part of $A$. Reactivity is thus equivalent to the maximum of the so-called Rayleigh quotient (Neubert & Caswell, 1997). This largest eigenvalue is precisely what is known as the epidemicity index in epidemiological contexts, highlighting the equivalence of these concepts. As noted by Townley et al. (2007), this concept was termed "*initial growth rate*" in Hinrichsen and Pritchard (2005). Reactivity has units of inverse time, $R_J > 0$ indicates that at least one perturbation initially grows, rendering the equilibrium reactive. For a given Jacobain of (3), $J$, and corresponding system (8), the set

$$X_J := \{x \in \mathbb{R}^n_+ : x(0) = x, xH(J)x^T > 0\},$$

has been termed the g-reactivity basin (Mari et al., 2017). Initial conditions for (8) in the g-reactivity basin, $X_J$, result in trajectories that have transient outbreaks. The magnitude of $R_J$ quantifies the potential extent of such short-term outbreaks. As noted in Hosack et al. (2008), by the Bendixson inequality, one can show that $\mu(J) \le \mu(H(J))$. So, in general, a non-reactive system has a stable disease-free equilibrium.

As with $J$, computing the eigenvalues of $H(J)$ is quite intractable. Similar to the role of $R_0$, we can derive a quantity, called a threshold index for epidemicity, that tells us if our system is reactive or not. To derive the threhold index for epidemicity we follow the derivation of Hosack et al. (2008). We can write $H(J) = H(J_F) + H(J_V)$. Since $J_V$ is diagonal, we have that $H(J_V) = J_V$. Therefore, $H(J_V)^{-1} = J_V^{-1}$ . We can compute that

$$H(J_F) = \frac{1}{2} \begin{pmatrix} 0 & \cdots & 0 & \dfrac{d_1 w_1 P_1 + V_T \nu_1 c_1}{P} \\ \vdots & \ddots & \vdots & \vdots \\ 0 & \cdots & 0 & \dfrac{w_n d_n P_n + V_T \nu_n c_n}{P} \\ \dfrac{d_1 w_1 P_1 + V_T \nu_1 c_1}{P} & \cdots & \dfrac{d_n w_n P_n + V_T \nu_n c_n}{P} & 0 \end{pmatrix}.$$

Therefore, we get that the so-called maximum next-generation matrix is

$$H(J_F)H(J_V)^{-1} = \frac{1}{2}\begin{pmatrix} 0 & \cdots & 0 & \dfrac{d_1 w_1 P_1 + V_T \nu_1 c_1}{\mu_V P} \\ \vdots & \ddots & \vdots & \dfrac{d_2 w_2 P_2 + V_T \nu_2 c_2}{\mu_V P} \\ 0 & \cdots & 0 & \dfrac{d_n w_n P_n + V_T \nu_n c_n}{\mu_V P} \\ \dfrac{d_1 w_1 P_1 + V_T \nu_1 c_1}{\mu_1 P} & \cdots & \dfrac{d_n w_n P_n + V_T \nu_n c_n}{\mu_n P} & 0 \end{pmatrix}, \tag{10}$$

and so the threshold index for epidemicity can be computed as

$$\mathcal{E}_0 := s\left(H(J_F)H(J_V)^{-1}\right) = \sqrt{\frac{1}{4P^2\mu_V}\sum_i \frac{(d_i w_i P_i + c_i \nu_i V_T)^2}{\mu_i}}.$$

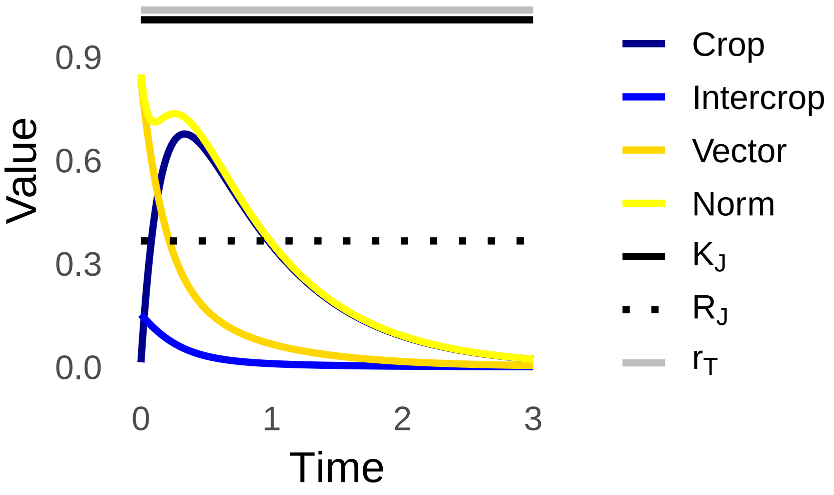


Fig. 4: Reactive and transient outbreak: Example of linearised dynamics of (3), showing trajectories $P_{1,I}$ (dark blue), $P_{2,1}$ (blue), $V_I$ (dark yellow) and the Euclidean norm of these trajectories (yellow) near the disease-free equilibrium. Also shown are the transient metrics $R_J > 0$ (black dotted), $K_J$ (black solid), and $r_T$ (grey solid). Parameters: $d_1 = 8.4, d_2 = 3.35, P_1 = 1000, P_2 = 200, \nu_1 = 1.88, \nu_2 = 1.96, w_1 = w_2 = 1, V_T = 50, c_1 = 6.54, c_2 = 12.14, \mu_1 = 2.63, \mu_2 = 5.93, \mu_V = 4.44$. Initial conditions: $P_{1,I}(0) = 0.013, P_{2,I}(0) = 0.152, V_I(0) = 0.835$.

#### 6.2 Amplification Envelope

The amplification envelope, defined as

$$r(t) = \|\exp(Jt)\|_2,$$

where $\exp(Jt)$ is the matrix exponential (O'Regan et al., 2020). This quantity measures the largest possible amplification of any unit-norm initial perturbation at time $t$. In practice, $r(t)$ corresponds to the largest singular value of $e^{Jt}$ and is computed numerically (O'Regan et al., 2020). Over a finite time interval $[0, T]$, we define the maximum amplification envelope over this interval as

$$r_T = \max_{t\in[0,T]} r(t).$$

Related to the maximum amplification envelope is the Kreiss bound, which is defined by

$$K_J = \sup_{\Re(z)>0} \left\{ \Re(z) \|(zI - J)^{-1}\|_2 \right\},$$

where $\Re(z)$ is the real part of $z \in \mathbb{C}$. This offers a worst-case estimate of transient amplification due to the non-normality of $J$ (Townley et al., 2007). The Kreiss Matrix Theorem ensures that

$$K_J \leq \sup_{t\geq 0} r(t) \leq \beta K_J,$$

where $\beta = \exp(1)n$ depends only on the system dimension $n$ (Raouafi, 2018). This inequality implies that, regardless of time, the amplification envelope $r(t)$ remains bounded between $K_J$ and $\beta K_J$. These bounds are conservative but provide a time-invariant envelope for the worst-case transient growth, even in systems that eventually return to equilibrium.

To illustrate transient dynamics, we simulated the linearised system of (3) for $n = 2$ (see Fig. 4). Although the infected compartments ultimately decline to extinction since $R_0 < 1$, the positive $R_J$ indicates reactivity. Therefore transient outbreaks are possible. For the initial condition chosen, we can see that a transient outbreak is realised. We can observe initial short-term growth in crop infection density, while other compartments show no amplification. The fact that transients remain bounded by $r_T$ and below the Kreiss bound highlights the value of transient metrics for predicting short-term outbreak severity, providing meaningful thresholds in systems that have a locally stable disease-free equilibrium, yet are reactive.

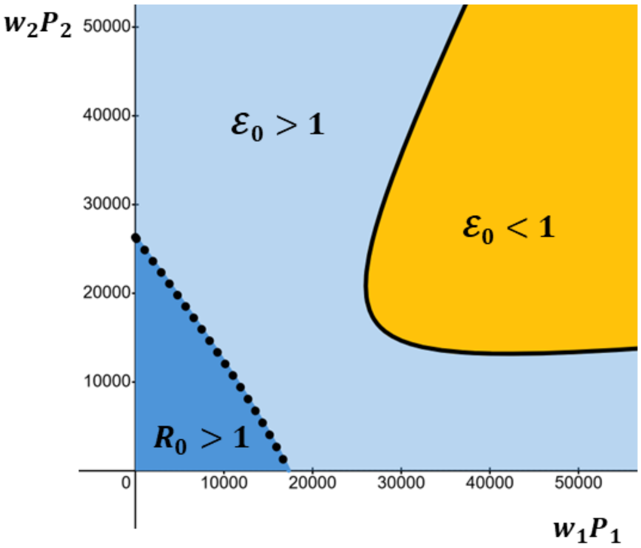


Fig. 5: Illustrative example of the $(w_1P_1, w_2P_2)$-plane where $\mathcal{E}_0 = 1$ does not intersect the positive coordinate axes. We can observe a reactive region (light blue), an initially resilient region (yellow), and a region where persistent outbreaks occur (dark blue). The solid black curve is $\mathcal{E}_0 = 1$ and the dotted black curve is $R_0 = 1$. Behaviour like this can be observed if one lets $c_1 = 4.1$, $c_2 = 2.8$, $d_1 = 8.6$, $d_2 = 4.9$, $\nu_1 = 1.7$, $\nu_2 = 1.4$, $\mu_1 = 6.3$, $\mu_2 = 1.4$, $V_T = 4920$ and $\mu_V = 2.5$.

### 7 Total Weighted Plant Densities and the Threshold Index for Epidemicity

In the context of our model for when $n = 2$, we can succintly summarise the behaviour of where transient and persistent outbreaks can occur in the $(w_1P_1, w_2P_2)$-plane by looking at the expressions for $R_0 = 1$ and $\mathcal{E}_0 = 1$, the proof of such behaviour will be given in Section 8).

1. If

$$4\mu_V > \frac{d_1^2 d_2^2}{d_1^2 \mu_2 + d_2^2 \mu_1} \tag{11}$$

then there exists one region where $\mathcal{E}_0 > 1$ (reactive), one region where $\mathcal{E}_0 < 1$ (initial resilience) and one region where $R_0 > 1$ (persistent outbreak). In this scenario there are different sub-cases, depending on if the region of initially resilience intersects the positive coordinate axes. We show in Section 8.4 that $\mathcal{E}_0 = 1$ crosses the positive $w_i P_i$-axis precisely when

$$d_i^2 < 4\mu_i \mu_V. \tag{12}$$

Alternatively, $\mathcal{E}_0 = 1$ does not cross the positive $w_i P_i$-axis when (12) does not hold. In Fig. 5 we can see an example where $\mathcal{E}_0 = 1$ does not intersect either coordinate axis. In Fig. 6 we can see an example where $\mathcal{E}_0 = 1$ intersects one of the axes. In Fig. 7 we can see an example where $\mathcal{E}_0 = 1$ intersects both axes. In the special case when the curves $R_0 = 1$ and $\mathcal{E}_0 = 1$ intersect at $(\hat{P}_1, \hat{P}_2)$ (see Section 8), there exists two regions where $\mathcal{E}_0 > 1$ (reactive), one region where $\mathcal{E}_0 < 1$ (initial resilience) and one region where $R_0 = 1$ (persistent outbreak). In Fig. 5-7, the yellow region is the region where $\mathcal{E}_0 < 1$, the dark blue region is where $R_0 > 1$, and the light blue region is where $\mathcal{E}_0 > 1$.
2. If (11) doesn't hold then there exists an unbounded region where $\mathcal{E}_0 > 1$ (reactive) and a finite region where $R_0 > 1$ (persistent outbreak) (see Fig. 8 for an illustrative diagram). In this case there is no region of initial resilience. In Fig. 8, the dark blue region is where $R_0 > 1$, and the light blue region is where $\mathcal{E}_0 > 1$.

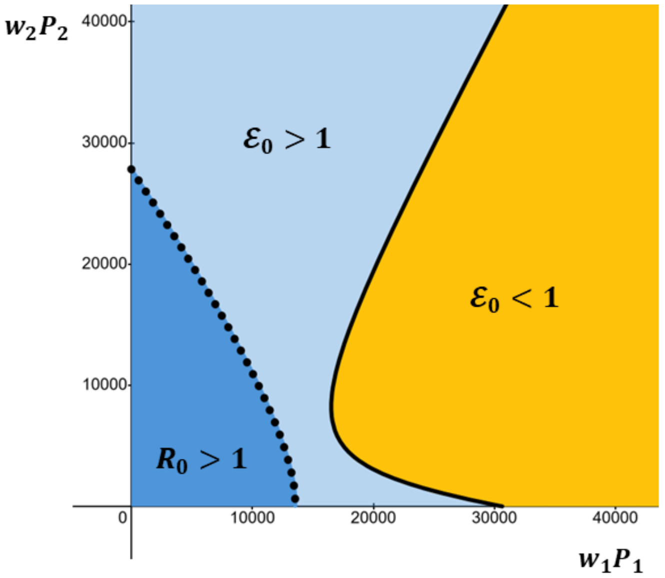


Fig. 6: Illustrative example of the $(w_1P_1, w_2P_2)$-plane where we can observe a reactive region (light blue), an initially resilient region (yellow) that intersects one of the positive coordinate axes, and a region where persistent outbreaks occur (dark blue). The solid black curve is $\mathcal{E}_0 = 1$ and the dotted black curve is $R_0 = 1$. Behaviour like this can be observed if one lets $c_1 = 8$, $c_2 = 3.6$, $d_1 = 8.6$, $d_2 = 5.5$, $\nu_1 = 1$, $\nu_2 = 1$, $\mu_1 = 10$, $\mu_2 = 1.4$, $V_T = 4920$ and $\mu_V = 2.5$.

We will conclude this section by briefly looking at two biologically motivated examples.

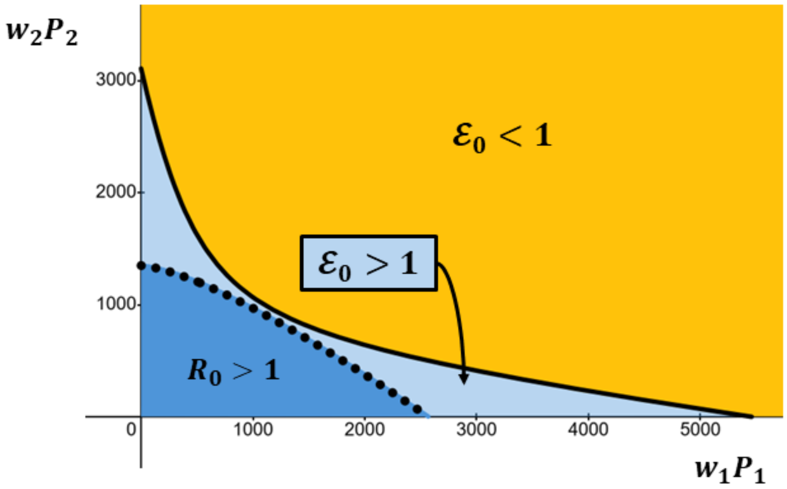


Fig. 7: Illustrative example of the $(w_1P_1, w_2P_2)$-plane where we can observe a reactive region (light blue), an initially resilient region (yellow) that intersects both positive axes, and a region where persistent outbreaks occur (dark blue). The solid black curve is $\mathcal{E}_0 = 1$ and the dotted black curve is $R_0 = 1$. Behaviour like this can be observed if one lets $c_1 = 2.5$, $c_2 = 1$, $d_1 = 7.2$, $d_2 = 6.3$, $\nu_1 = 0.5$, $\nu_2 = 0.6$, $\mu_1 = 7$, $\mu_2 = 5.6$, $V_T = 5000$ and $\mu_V = 2.5$.

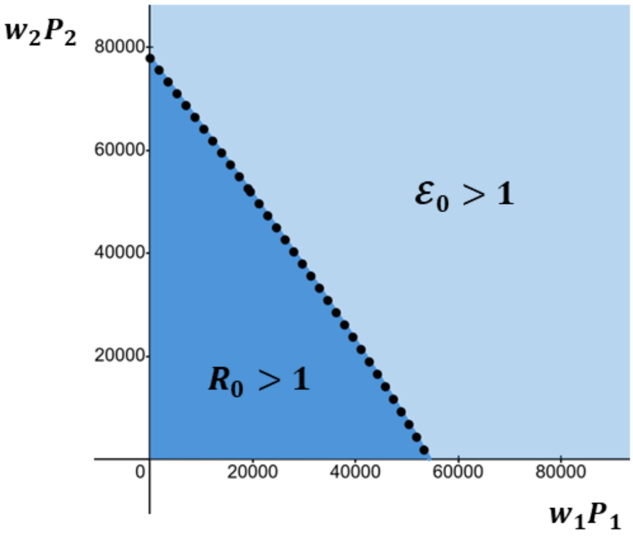


Fig. 8: Illustrative example of the $(w_1P_1, w_2P_2)$-plane where we can observe only a reactive region (light blue) and a region where persistent outbreaks occur (dark blue). The dotted black curve is $R_0 = 1$. Parameters: $c_1 = 4$, $c_2 = 3.6$, $\nu_1 = 2$, $\nu_2 = 1$, $d_1 = 7.6$, $d_2 = 7.7$, $\mu_1 = 2.2$, $\mu_2 = 0.7$, $V_T = 4920$ and $\mu_V = 2.5$.

#### 7.1 Example 1: Identical Transmission

Assume $p_1 = p_2 = p > 0$ for any parameter $p_i \in \{c_i, \nu_i, d_i, \mu_i\}$. In this neutral scenario, the only difference in the two plant hosts is the total weighted abundances/densities $w_1P_1$ and $w_2P_2$. Therefore we get that

$$R_0 = \sqrt{\frac{V_T c \nu d}{(w_1P_1 + w_2P_2)\mu\mu_V}}.$$

We can see that $R_0 = 1$ is always a straight line in the $(w_1P_1, w_2P_2)$-plane (see Fig. 9). We also get that

$$\mathcal{E}_0 = \sqrt{\frac{(dw_1P_1 + c\nu V_T)^2 + (dw_2P_2 + c\nu V_T)^2}{4P^2\mu_V\mu}}.$$

Although $\mathcal{E}_0 = 1$ is not a straight line in the $(w_1P_1, w_2P_2)$-plane, due to the quadratic terms, we do have symmetry of the curve $\mathcal{E}_0 = 1$ when you reflect about the line $w_1P_1 = w_2P_2$. This is because it is invariant if you swap $w_1P_1$ with $w_2P_2$.

This scenario is a special case of the general result derived in Section 8.4. If we let $A = c\nu V_T/d$ and $B = d^2/4\mu\mu_V$, we can conclude that when $B < 1$ and an initial resilience region exists, it intersects both positive coordinate axes at the same point. This point can be computed as

$$\hat{P} = \frac{-BA - A\sqrt{B(2-B)}}{B-1}. \tag{13}$$

We can visualise what this behaviour looks like in Fig. 9. It is intuitive that in a monculture setting, if $B < 1$ ($d$ is relatively small, for example) both crop and intercrop systems enter a region of initial resilience for the total weighted plant density above $\hat{P}$. Once these crops are intercropped, there are weighted mixtures of each where they exhibit initial resilience for lower $P_1$ and $P_2$ values. Essentially all one is doing is increasing $P = w_1P_1 + w_2P_2$ beyond $\hat{P}$, as both hosts are identical (all parameters are equal).

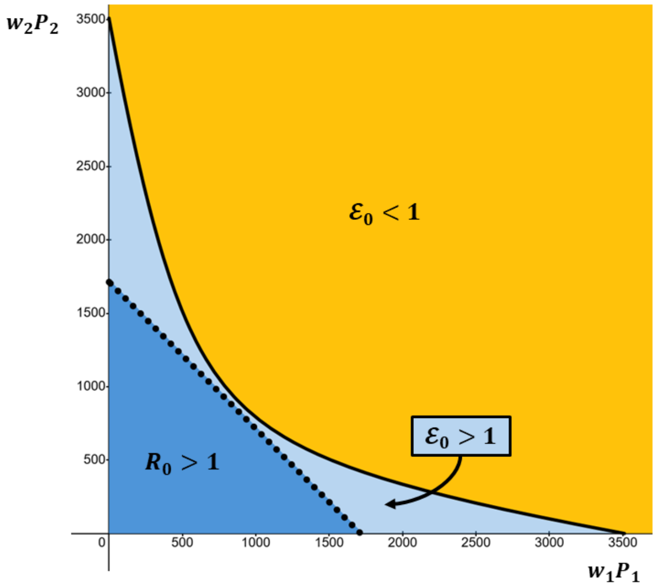


Fig. 9: Illustrative example of the $(w_1P_1, w_2P_2)$-plane for the “neutral” example, when the region of initial resilience intersects both positive coordinate axes at $\hat{P} \approx 3500$. We also observe a reactive region (light blue) and a region where persistent outbreaks occur (dark blue). The dotted black curve is $R_0 = 1$. Parameters: $c = 0.6$, $\nu = 1$, $d = 10$, $\mu = 3.5$, $V_T = 10{,}000$ and $\mu_V = 10$.

### 7.2 Example 2: Varietal Mixtures

Assume that an initially resilient region exists, i.e. (11) holds. Further assume that both the crop and intercrop are varietal mixtures, where instead of being different species, they are varieties of the same crop (see main text for examples). In this case it would be interesting to look at the line $w_1P_1 + w_2P_2 = P$ in the $(w_1P_1, w_2P_2)$-plane, to see what constant total weighted plant population gives certain dynamical behaviour, for some chosen $P > 0$. In Fig. 10 we can see an example of this for when, for example, initial resilience cannot be observed for both the crop and the intercrop monoculture system, i.e. $B_1 < 1$ and $B_2 < 1$ (see Section 8.4 for details).

In this example, the regime one enters depends on the choice of $P > 0$, which in turn is both context- and species-dependent. We show three lines for three different $P$ values to illustrate a few of the many scenarios one can observe. In the first case (solid red line) all weighted varietal mixture combinations result in persistent outbreaks. This could be due to concentration effects. In the second case (dashed red line) all weighted varietal mixture combinations result the potential for transient outbreaks. In the final case (dotted red line) no persistent outbreaks can occur, but for both large enough weighted crop/intercrop densities transient outbreaks may occur. In this scenario there are intermediate weighted varietal mixture combinations that allow for initial resilience to emerge, once there is a sufficient numbers of both crops and intercrops to help tip $\mathcal{E}_0$ below 1.

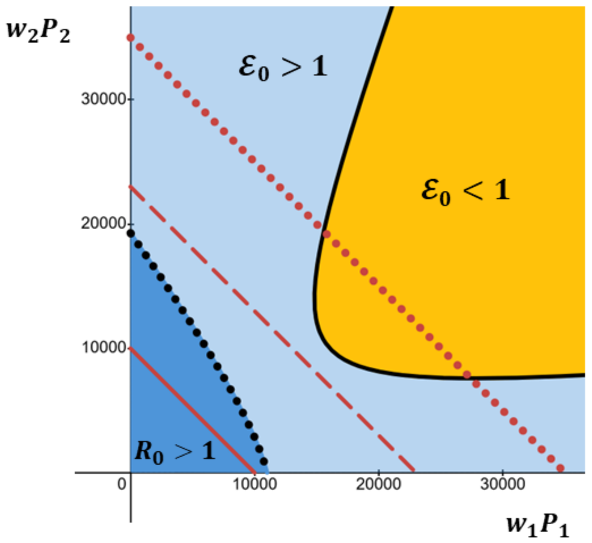


Fig. 10: Illustrative example of the $(w_1P_1, w_2P_2)$-plane where $\mathcal{E}_0 = 1$ does not intersect the positive coordinate axes. We can observe a reactive region (light blue), an initially resilient region (yellow), and a region where persistent outbreaks occur (dark blue). The solid black curve is $\mathcal{E}_0 = 1$ and the dotted black curve is $R_0 = 1$. The red lines represent the curve $w_1P_1 + w_2P_2 = P$ for three different $P > 0$ values. Parameters: $c_1 = 4.1$, $c_2 = 2.8$, $\nu_1 = 1$, $\nu_2 = 1$, $d_1 = 8.6$, $d_2 = 4.9$, $\mu_1 = 6.3$, $\mu_2 = 1.4$, $V_T = 4920$ and $\mu_V = 2.5$.

### 8 Behaviour of $R_0 = 1$ and $\mathcal{E}_0 = 1$ while Varying $w_1P_1$ and $w_2P_2$

We will now describe the general behaviour of $R_0 = 1$ and $\mathcal{E}_0 = 1$ in the $(w_1P_1, w_2P_2)$-plane that we observed in Section 7. Let $x = w_1P_1$ and $y = w_2P_2$ denote total weighted crop and intercrop abundances, respectively. We use this notation here for ease of exposition. Also fix the parameters $c_i > 0$, $\nu_i > 0$, $d_i > 0$, $\mu_i > 0$, $\mu_V > 0$ for $i \in \{1, 2\}$. In order to characterise the behaviour of $R_0 = 1$ and $\mathcal{E}_0 = 1$ in the positive quadrant, we will rewrite $R_0 = 1$ as

$$\frac{c_1\nu_1 d_1 x}{\mu_1} + \frac{c_2\nu_1 d_2 y}{\mu_2} = \frac{\mu_V (x+y)^2}{V_T}$$

rewrite $\mathcal{E}_0 = 1$ as

$$\frac{(d_1 x + \nu_1 c_1 V_T)^2}{\mu_V \mu_1} + \frac{(d_2 y + \nu_2 c_2 V_T)^2}{\mu_V \mu_2} = 4(x+y)^2.$$

This is so we can look at $x$ and $y$ values in $\mathbb{R}^2_+$ and $\mathbb{R}^2_-$. This will help us definitively show how these curves behave in the positive quadrant.

#### 8.1 $R_0 = 1$ and $\mathcal{E}_0 = 1$ can only intersect at a single point in $\mathrm{Int}(\mathbb{R}^2_+)$

Setting $R_0 = 1$ and $\mathcal{E}_0 = 1$ equal gives

$$\frac{d_1^2\left(\frac{\nu_1 c_1 V_T}{d_1} + x\right)^2}{\mu_1} + \frac{d_2^2\left(\frac{\nu_2 c_2 V_T}{d_2} + y\right)^2}{\mu_2} = 4V_T\left(\frac{\nu_1 c_1 d_1 x}{\mu_1} + \frac{\nu_2 c_2 d_2 y}{\mu_2}\right).$$

This is quadratic in $y$, with discriminant

$$\Delta = \frac{-4d_2^2(d_1 x - \nu_1 c_1 V_T)^2}{\mu_1 \mu_2}.$$

We need there to exist a real solution of this quadratic. This occurs only if

$$d_1 x - \nu_1 c_1 V_T = 0 \iff x = \frac{\nu_1 c_1 V_T}{d_1}.$$

By an analagous argument using a quadratic in $x$, we can conclude that the only intersection of $R_0 = 1$ and $\mathcal{E}_0 = 1$ occurs at

$$(\hat{P}_1, \hat{P}_2) := \left(\frac{\nu_1 c_1 V_T}{d_1}, \frac{\nu_2 c_2 V_T}{d_2}\right) \in \mathrm{Int}(\mathbb{R}^2_+).$$

#### 8.2 $R_0 = 1$ is always a parabola that intersects $(0, 0)$ and $\mathrm{Int}(\mathbb{R}^2_+)$

Viewed as a conic section, the curve $R_0 = 1$ can be written as

$$\frac{\mu_V}{V_T}x^2 + \frac{2\mu_V xy}{V_T} + \frac{\mu_V y^2}{V_T} - \frac{\nu_1 c_1 d_1}{\mu_1}x - \frac{\nu_2 c_2 d_2}{\mu_2}y = 0.$$

The discriminant of this quadratic is $\Delta = 0$, so the curve is always a parabola. This curve intersects the $x$-axis at 0 and

$$\frac{\nu_1 c_1 d_1 V_T}{\mu_1 \mu_V} > 0$$

and intersects the $y$-axis at 0 and

$$\frac{\nu_2 c_2 d_2 V_T}{\mu_2 \mu_V} > 0.$$

#### 8.3 $\mathcal{E}_0 = 1$ is always a hyperbola when it intersects $\mathrm{Int}(\mathbb{R}^2_+)$

We will now show that when $\mathcal{E}_0 = 1$ intersects the positive quadrant it is a hyperbola. In particular, we will show that one and only one branch of this hyperbola intersects the positive quadrant.

For a fixed $P \in \mathbb{R}$, consider the line

$$x + y = P. \tag{14}$$

Also consider the ellipse

$$B_1(A_1 + x)^2 + B_2(A_2 + y)^2 = P^2, \tag{15}$$

centred at $(-A_1, -A_2)$, where

$$A_i = \frac{\nu_i c_i}{d_i}V_T, \quad B_i = \frac{d_i^2}{4\mu_i \mu_V}.$$

One can confirm that a point $(x, y) \in \mathbb{R}^2$ lies on the curve $\mathcal{E}_0 = 1$ if and only if it simultaneously satisfies (14) and (15) for some chosen $P$. Therefore, the intersections of (14) and (15) generates all points of the curve $\mathcal{E}_0 = 1$ as $P$ varies from 0 to $\infty$. Intersections of (15) with the line $x + y = P$ for $P < 0$ lie entirely below the line $y = -x$, and so these intersections are not biologially relevant for our model.

For the line $x + y = P$, with $P > 0$, substituting $y = P - x$ into (15) gives a quadratic in $x$

$$(B_1 + B_2)x^2 + 2(B_1A_1 - B_2(A_2 + P))x + B_1A_1^2 + B_2(A_2 + P)^2 - P^2 = 0. \tag{16}$$

Assume that this quadratic has two real solutions. This is equivalent to assuming that

$$\Delta = (B_1 + B_2)P^2 - B_1B_2(A_1 + A_2 + P)^2 > 0 \iff \frac{B_1 + B_2}{B_1B_2} > \left(\frac{A_1 + A_2}{P} + 1\right)^2.$$

Since this last inequality holds, and we know $A_i > 0$ and $P > 0$, we then have that

$$\left(\frac{A_1 + A_2}{P} + 1\right)^2 > 1 \implies \frac{B_1 + B_2}{B_1B_2} > 1. \tag{17}$$

This last inequality in (17) can be written as

$$4\mu_V > \frac{d_1^2 d_2^2}{d_1^2\mu_2 + d_2^2\mu_1}. \tag{18}$$

We can rewrite the curve $\mathcal{E}_0 = 1$ in standard conic section form as

$$(B_1 - 1)x^2 - 2xy + (B_2 - 1)y^2 + 2B_1A_1x + 2B_2A_2y + B_1A_1^2 + B_2A_2^2 = 0.$$

This conic section is a hyperbola when it's discriminant is positive. One can confirm that occurs precisely when (18) holds. Since the intersections of (14) and (15) with $P > 0$ imply that (18) holds, we must therefore have that one branch of the hyperbola $\mathcal{E}_0 = 1$ satisfies $x + y > 0$, so this branch lies entirely outside the negative quadrant and intersects the positive quadrant. This must imply then that the other branch is below the line $y = -x$, and so it lies entirely outside the positive quadrant and intersects the negative quadrant.

Note that if (18) does not hold, we get that the conic section $\mathcal{E}_0 = 1$ can be either a parabola or an ellipse. In these two cases, we have that (16) has no positive real solutions for any $P > 0$, and so the curve $\mathcal{E}_0 = 1$ does not intersect the positive quadrant.

We conjecture that the arguments above generalise to the case when $n \geq 2$, with the harmonic mean condition (18) becoming

$$4\mu_V > \left(\sum_{i=1}^{n} \frac{\mu_i}{d_i^2}\right)^{-1}. \tag{19}$$

In this case one would have to prove (19) is sufficient to observe similar behaviour within the $n$-dimensional space $w_1P_1 \times \cdots \times w_nP_n$, using the theory of quadrics and hyperplanes.

#### 8.4 If $\mathcal{E}_0 = 1$ is a hyperbola it intersects the positive $w_iP_i$-axis when $d_i^2 < 4\mu_i\mu_V$.

We will now demonstrate how $B_i < 1$ or $B_i \geq 1$ determines when the curve $\mathcal{E}_0 = 1$, assumed to be a hyperbola, intersects the positive $w_iP_i$-axis.

Fix $A_1, A_2, B_1, B_2 > 0$. From (15) and (14), we can see that points on the $\mathcal{E}_0 = 1$ curve lying on the positive $x$-axis correspond to values of $P$ where

$$B_1(A_1 + x)^2 + B_2A_2^2 = P^2 \tag{20}$$

and $x = P$ coincide. Let

$$z_1(P) := -A_1 + \sqrt{\frac{P^2 - B_2 A_2^2}{B_1}},$$

defined for $P \geq \sqrt{B_2}A_2$, and

$$z_2(P) := P.$$

For a given $P$, $z_1(P)$ is the value of $x$ such that (20) holds and $z_2(P)$ is the value of $x$ such that $x = P$. Hence, if $z_1$ and $z_2$ intersect, then $\mathcal{E}_0 = 1$ intersects the $x$-axis. Note that both $z_1$ and $z_2$ are continuous and strictly increasing functions of $P > \sqrt{B_2}A_2$.

At the smallest feasible $P = P_{min} := \sqrt{B_2}A_2$, we have that

$$z_1(P_{min}) = -A_1 < 0, \qquad z_2(P_{min}) = \sqrt{B_2}A_2 > 0.$$

Therefore $z_1(P) < z_2(P)$ for $P$ in a neighbourhood of $P_{min}$.

We have three cases to consider:

1. Assume $B_1 > 1$, then we can see that $1/\sqrt{B_1} < 1$ and so

$$z_1(P) < \sqrt{\frac{P^2 - B_2 A_2^2}{B_1}} \leq \frac{P}{\sqrt{B_1}} < z_2(P).$$

   Hence $z_1(P) < z_2(P)$ for all $P \geq \sqrt{B_2}A_2$, and no solution of $z_1(P) = z_2(P)$ exists. The curve therefore never intersects the $x$-axis.
2. Assume $B_1 < 1$. Notice that

$$f(P) := z_1(P) - z_2(P) = -A_i + \frac{P}{\sqrt{B_1}}\sqrt{1 - \frac{B_2 A_2^2}{P^2}} - P$$

   for $P \geq \sqrt{B_2}A_2$. For sufficiently large $P$ we can see that

$$f(P) \approx -A_i + \sqrt{P}\left(\frac{1}{\sqrt{B_1}} - 1\right). \tag{21}$$

   Since $B_1 < 1$, and so $\sqrt{B_1} < 1$, then we can see from (21) that there must exist some $P' > 0$ sufficiently large, such that $f(P') > 0$. The function $f$ is continuous on $[\sqrt{B_2}A_2, \infty)$. Therefore, since $f(P_{min}) < 0$ and $f(P') > 0$, by the Intermediate Value Theorem, there exists $\hat{P} \in (P_{min}, P')$ such that

$$f(\hat{P}) = 0 \implies z_1(\hat{P}) = z_2(\hat{P}).$$

   The curve $\mathcal{E}_0 = 1$ therefore intersects the $x$-axis at $\hat{P} > 0$.
3. We can write $\mathcal{E}_0 = 1$, with $y = 0$, as

$$(B_1 - 1)x^2 + 2B_1 A_1 x + B_1 A_1^2 + B_2 A_2^2 = 0. \tag{22}$$

   Assume $B_1 = 1$. We can see then that

$$x = \frac{-A_1^2 - B_2 A_2^2}{2} < 0,$$

   and so $\mathcal{E}_0 = 1$ cannot intersect the postive $x$-axis.

Analogous arguments to those above apply when deriving conditions for when the curve $\mathcal{E}_0 = 1$ intersects the $y$-axis. We also conjecture that these arguments can be generalised to the case when $n \geq 2$.
